# The Neurolipid Atlas: a lipidomics resource for neurodegenerative diseases uncovers cholesterol as a regulator of astrocyte reactivity impaired by ApoE4

**DOI:** 10.1101/2024.07.01.601474

**Authors:** Femke M. Feringa, Sascha J. Koppes-den Hertog, Lian Wang, Rico J.E. Derks, Iris Kruijff, Lena Erlebach, Jorin Heijneman, Ricardo Miramontes, Nadine Pömpner, Niek Blomberg, Damien Olivier-Jimenez, Lill Eva Johansen, Alexander J. Cammack, Ashling Giblin, Christina E Toomey, Indigo V.L. Rose, Hebao Yuan, Michael Ward, Adrian M. Isaacs, Martin Kampmann, Deborah Kronenberg-Versteeg, Tammaryn Lashley, Leslie M. Thompson, Alessandro Ori, Yassene Mohammed, Martin Giera, Rik van der Kant

**Affiliations:** Center for Neurogenomics and Cognitive Research, Vrije Universiteit Amsterdam, Amsterdam Neuroscience, Amsterdam, the Netherlands; Alzheimer Center Amsterdam, Department of Neurology, Amsterdam University Medical Center, Amsterdam Neuroscience, Amsterdam, the Netherlands; Leiden University Medical Center, Center for Proteomics and Metabolomics, Leiden, the Netherlands; German Center for Neurodegenerative Diseases (DZNE) Tübingen, Tübingen, Germany; Department of Cellular Neurology, Hertie Institute for Clinical Brain Research, University of Tübingen, Tübingen, Germany; Department of Psychiatry and Human Behavior, University of California, Irvine, CA, USA; Department of Neurobiology and Behavior, University of California, Irvine, CA, USA; Institute for Memory Impairments and Neurological Disorders, University of California, Irvine, CA, USA; Leibniz Institute on Aging, Fritz Lipmann Institute, Jena, Germany; Department of Neurodegenerative Disease, UCL Queen Square Institute of Neurology, University College London, London, UK; UK Dementia Research Institute at UCL, University College London, London, UK; Department of Clinical and Molecular Neuroscience, Queen Square Institute of Neurology, University College London, London, UK; Institute for Neurodegenerative Diseases and Neuroscience Graduate Program, University of California, San Francisco, San Francisco, CA, USA; National Institute of Neurological Disorders and Stroke, National Institutes of Health, Bethesda, MD, USA; Department of Biochemistry and Biophysics, Institute for Neurodegenerative Diseases, University of California, San Francisco, San Francisco, CA, USA; Gerald Bronfman Department of Oncology, McGill University, Montreal, QC H3A 0G4, Canada

**Author notes:** co-first (Femke M. Feringa, Sascha J. Koppes-den Hertog).

## Abstract

Lipid changes in the brain have been implicated in many neurodegenerative diseases including Alzheimer’s Disease (AD), Parkinson’s disease and Amyotrophic Lateral Sclerosis. To facilitate comparative lipidomic research across brain-diseases we established a data commons named the Neurolipid Atlas, that we have pre-populated with novel human, mouse and isogenic induced pluripotent stem cell (iPSC)-derived lipidomics data for different brain diseases. We show that iPSC-derived neurons, microglia and astrocytes display distinct lipid profiles that recapitulate *in vivo* lipotypes. Leveraging multiple datasets, we show that the AD risk gene ApoE4 drives cholesterol ester (CE) accumulation in human astrocytes recapitulating CE accumulation measured in the human AD brain. Multi-omic interrogation of iPSC-derived astrocytes revealed that cholesterol plays a major role in astrocyte interferon-dependent pathways such as the immunoproteasome and major histocompatibility complex (MHC) class I antigen presentation. We show that through enhanced cholesterol esterification ApoE4 suppresses immune activation of astrocytes. Our novel data commons, available at neurolipidatlas.com, provides a user-friendly tool and knowledge base for a better understanding of lipid dyshomeostasis in neurodegenerative diseases.

## Introduction

As one of the most lipid rich organs in our body^1^, the brain heavily relies on proper brain lipid homeostasis. Mutations in lipid metabolic genes cause rare, but severe, juvenile neurodegenerative diseases such as neuronal ceroid lipofuscinoses^2^ and Niemann Pick Type C^3^. More recently, changes in lipid metabolism have been implicated in common neurodegenerative diseases such as Alzheimer’s Disease (AD)^4–10^, Parkinson’s Disease (PD)^11,12^, Huntington’s Disease^13–15^, spinocerebellar ataxia^16^, Amyotrophic Lateral Sclerosis (ALS)^17,18^ and frontotemporal dementias including primary tauopathies^19–22^. In addition, conditions associated with neurodegenerative disease pathogenesis such as aging^23^, microglial activation by demyelination or fibrillar amyloid beta^24–26^, astrocyte activation^27^ or even altered sleep cycles^28^ have been recently shown to dysregulate brain lipid metabolism.

Together these findings strongly indicate that alterations in brain lipid metabolism can contribute to neurodegenerative diseases. More importantly, these findings suggest that lipid-targeting interventions could be a promising therapeutic strategy to prevent or treat these diseases. The exact number of endogenous mammalian lipids is unknown. However, it is likely that thousands of individual lipid species together shape cell specific lipidomes (lipotypes) that dictate cellular function and dysfunction in the brain^29^. Yet, sufficient detail on the exact lipid species and downstream pathways that contribute to the different neurodegenerative diseases is lacking. Mapping the primary disease-associated changes in the human brain lipidome is especially challenging, as confounders such as aging, diet, post-mortem interval, and secondary neurodegenerative processes (e.g. cell death) all strongly affect lipid metabolism. While animal models have been instrumental for our progress in understanding neurodegeneration, they have limited use for the study of lipids, as the human brain lipidome is intrinsically more complex^30^. Furthermore, studies of lipid metabolism in the human or rodent brain are typically performed in bulk brain tissue, not capturing cell-type specific changes as for example in neurons, astrocytes and microglia. A potential solution to overcome these challenges is induced pluripotent stem cell (iPSC) technology. Especially in combination with CRISPR/Cas9 gene-editing, this technology provides a powerful tool to study how disease-specific mutations and risk variants affect downstream disease phenotypes (e.g. Amyloid overproduction, pTau levels, αSynuclein levels)^31–34^. Furthermore, iPSC-models are scalable, allowing high-throughput drug discovery^7^.

To understand genotype-lipid interactions in human brain cells, here we have developed a standardized pipeline that combines isogenic iPSC-technology and lipidomics analysis capable of quantifying more than 1000 different lipid species. We also generated a lipidomics data commons, the Neurolipid Atlas (available at the www.neurolipidatlas.com) that allows for user-friendly exploration of (neuro)lipidomics data. We prepopulated the Neurolipid Atlas with data from a variety of different human iPSC-derived disease models and states (AD, PD, ALS, FTD), as well as post-mortem derived human and mouse brain as benchmarking datasets. Additionally, using this pipeline and data analysis tool, we show for the first time that iPSC-derived neurons, astrocytes and microglia have distinct lipid profiles resembling *in vivo* lipotypes. Through comparative lipidomic profiling of APOE3/3, APOE4/4 and reactive APOE3/3 iPSC-derived astrocytes, we show that cholesterol esters (CEs) and triacylglycerides (TGs) accumulate in ApoE4 iPSC-derived astrocytes (as in AD brain), but decrease in activated astrocytes. Through proteomic and functional characterization, we show that cholesterol metabolism directly controls astrocyte activation and interferon-dependent pathways such as the immunoproteasome and major histocompatibility complex (MHC) class I antigen presentation. High levels of free cholesterol enhance immune activation, whereas cholesterol esterification (increased in ApoE4 astrocytes) buffers immune activation. Overall, our Neurolipid Atlas provides a novel lipidomics tool and resource for the neuro field forming a cornerstone for future research into cell and (neurodegenerative) disease specific alterations of lipid metabolism. We exemplify the potential of our tool through generating proof of concept that altered cholesterol metabolism and CE accumulation in ApoE4 astrocytes hampers their immune function.

### Lipid profiles of human iPSC-derived neurons, astrocytes and microglia recapitulate known *in vivo* lipotypes

To allow easy exploration, analysis and sharing of brain lipidomics data, we generated a novel resource that we named Neurolipid Atlas (Fig 1A). This resource consists of two modules: one module containing datasets generated from iPSC-derived brain cells and one module containing newly generated data from human and mouse post-mortem brain samples (Fig 1A, discussed below). To populate the database, we developed a standardized iPSC-lipidomics pipeline capable of quantifying >1000 lipid species across 17 different classes in a cell-type specific manner (Fig 1A). iPSC-derived brain cells have robustly been shown to resemble *in vivo* brain cell-types at the transcript level (albeit more immature)^35–38^. Whether iPSC-derived neurons, astrocytes and microglia also resemble the *in vivo* lipidome is not known. Consequently, we differentiated iPSCs from a control iPSC line (BIONi037-A^39^) into glutamatergic neurons by Zhang *et al.* 2013^40^, astrocytes by Fong *et al.* 2018^41^ and microglia by Haenseler *et al.* 2017^35^ (Fig 1B) and confirmed cell fate with cell-type specific markers MAP2 (neurons), AQP4 (astrocytes) and Iba1 (microglia) (Fig 1C). We analyzed their lipidome by comprehensive, quantitative shotgun lipidomic analysis^42,43^ and found that iPSC-derived neurons, astrocytes and microglia had very distinct lipid profiles (Fig 1D-G, individual lipid species in Sup Fig 1) that resembled lipotypes of freshly isolated cells from mouse brain tissue^44^. Phosphatidylcholine (PC) and phosphatidylethanolamine (PE) were the most abundant lipid class in all cell types. Consistent with mouse brain cells we observed highest relative PC and PE levels in neurons^44^ (Fig 1D-G). Also, the PC and PE derivative lysophospholipids (LPC and LPE) were most abundant in neurons. Sphingomyelins (SM) were highly abundant in microglia, with lower levels in astrocytes and very low levels in neurons, similar to freshly isolated murine cells (Fig 1D,F-G)^44^. Also consistent with mouse data, phosphatidylserines (PS) were most abundant in microglia and astrocytes but low in neurons (Fig 1D,F-G), while neurons had the highest relative levels of ceramides (CER; Fig 1D-G)^44^. Diacylglyceride (DG) levels were highest in astrocytes, consistent with mouse data. Not in keeping with the mouse data were the relatively high DG levels in our iPSC-derived microglia, while phosphatidylglycerol (PG) lipids were relatively low (Fig 1G)^44^. Of the lipid classes that were not measured in the previous mouse study, we found that triacylglycerides (TG) and free fatty acids (FA) were most abundant in microglia. Astrocytes had the highest cholesterol ester (CE) stores, in line with the role of astrocytes as cholesterol supplier for other brain cell types^45,46^. Overall, these data indicate that iPSC-derived neurons, astrocytes and microglia not only recapitulate brain cells at the transcriptomic and proteomic level, but also at the lipidomic level. As for the remainder of the manuscript, all the lipidomics data is available through the Neurolipid Atlas (www.neurolipidatlas.com) where it can be explored, analyzed and downloaded.

**Figure 1.**
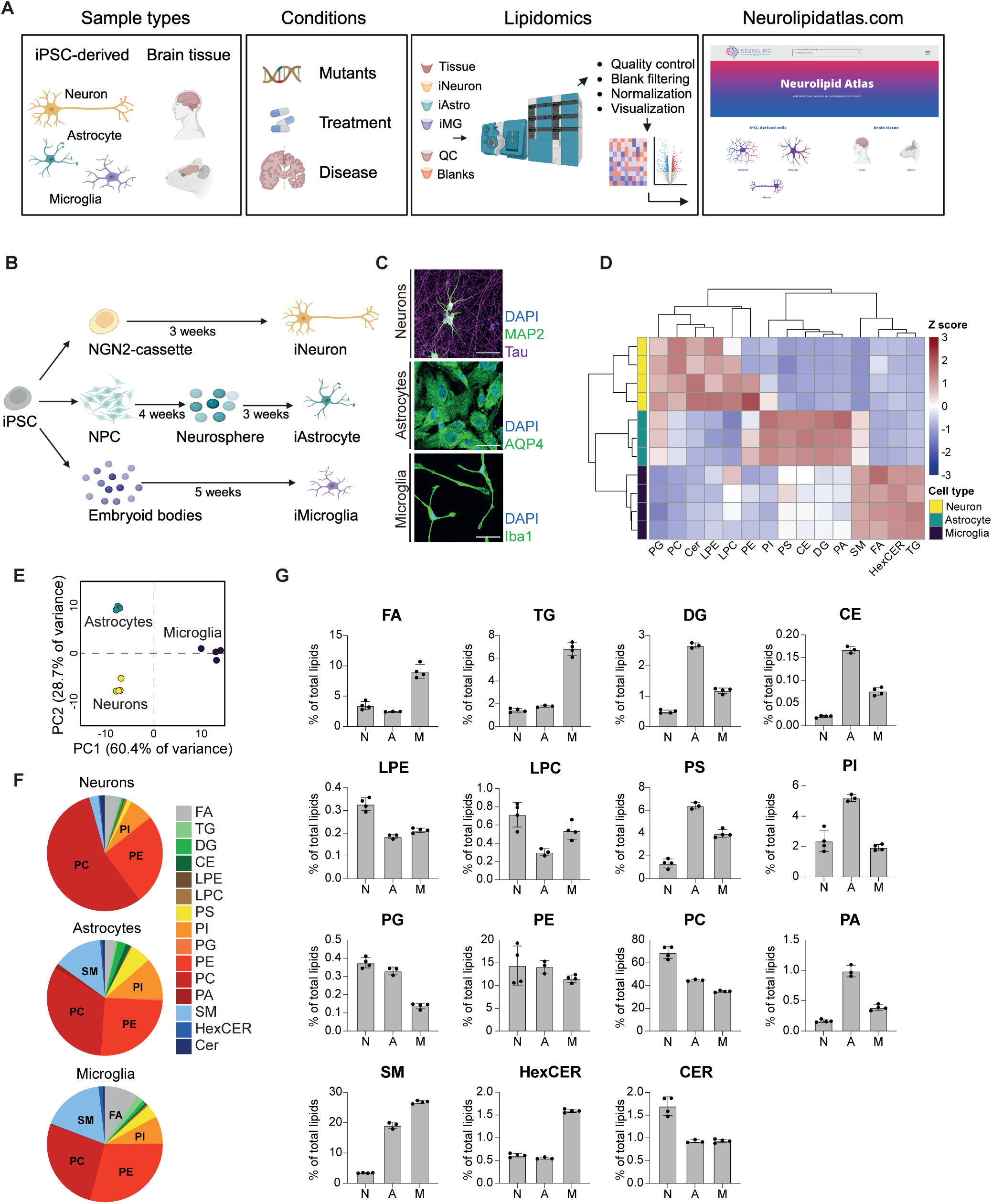
Lipotypes of human iPSC-derived neurons, astrocytes and microglia. A) Schematic overview of the Neurolipid Atlas work-flow and resource. B) Schematic overview of iPSC differentiation protocols. C) Representative confocal microscopy image of iPSC-derived neurons, astrocytes and microglia in monoculture. Scale bar = 50mm. D) Heatmap of Z scored lipid class abundance in iPSC-derived neurons, astrocytes and microglia (BIONi037-A parental line). E) PCA analysis of iPSC-derived brain cell lipotypes. F) Pie charts showing relative abundance of all detected lipid classes in the iPSC-derived brain cell types. G) Bar graphs present individual lipid class levels in each cell type, normalized to total lipid level. N (Neurons) n=4 wells, A (Astrocytes) n=3 wells, M (Microglia) n=4 wells. Mean + SD.

### Cholesterol esters accumulate in the human sporadic AD brain

The second module of the Neurolipid Atlas (Fig 1A) contains lipidomic data from human and mouse brain tissue. Here we focused on lipid changes in the AD brain. Only a handful of previous studies have performed lipidomic analysis on human AD brain tissue^4,47,48^. We determined the control (n=13) and AD (n=20) lipidome across three different brain areas (Fig 2A). We selected a brain area where AD pathology is abundant (frontal cortex) and an area where pathology is generally low or absent (cerebellum)^49^. In addition, within the frontal cortex we differentiated between gray matter (low in oligodendrocytes) and white matter (rich in oligodendrocytes). First, we explored regional differences in lipid composition between brain regions in the control brains (Fig 2B-C, Sup Fig 2A). The frontal cortex white matter had high levels of ceramides (CER, HexCER) and sphingomyelin (SM), consistent with the enrichment of these lipids in oligodendrocytes (Fig 2C, Sup Fig 2A)^44,50^. On the contrary, phospholipid and storage lipid (e.g. CE, TG) levels were relatively higher in cerebellum and frontal cortex gray matter (Fig 2C, Sup Fig 2A). Next, we explored differences in lipid composition between AD and control brains for each brain region. Principal component analysis (PCA) largely separated control and AD samples in the frontal cortex gray and white matter, but less so in the cerebellum (Fig 2D, Sup Fig 2B-C). At the class level, we found that cholesterol esters were significantly upregulated in AD in frontal cortex gray and white matter (2E-G, Sup Fig 2J-Q) and trended towards increased levels in cerebellum. Analysis at the level of individual lipid species also showed an increase for most CE species (sup 2D-F), but no single CE species reached significance, likely reflecting high variation in fatty acid tails of CEs in individual human subjects. In addition, TG levels (trend in all areas) and DG levels (significant only for frontal cortex white matter) were increased in AD subjects (Fig 2E-G). Lactosylceramides (LacCER) were also significantly increased in AD frontal cortex white matter (Fig 2E-F). Since astrogliosis is known to be increased in late stages of AD and reactive astrocytes adopt a distinct lipid profile with increased phospholipid saturation^27^, we looked at saturation of phospholipids and TGs in AD versus control brain tissue. However, no consistent changes in phospholipid or TG saturation were observed (Sup Fig 2G-I). Our results, combined with previous findings^4,47,48^ in AD subjects, strongly suggest that CE accumulation is a specific and key (lipidomic) feature of AD.

**Figure 2.**
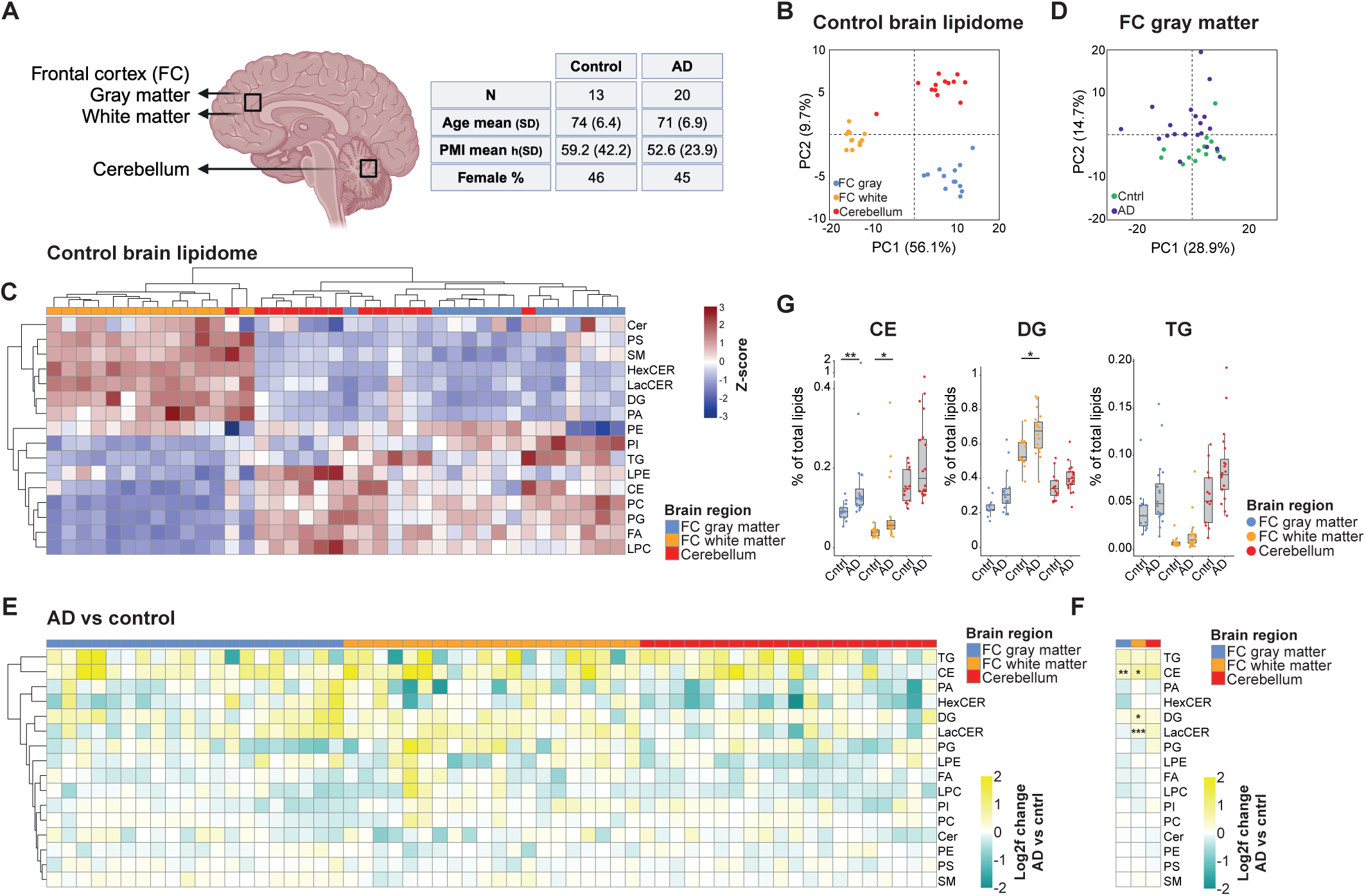
Human (AD) brain lipidomics. A) Schematic overview of human postmortem brain tissue sampled and summary of subject characteristics. Metadata for individual patients can be found in the methods section. B) PCA plot of unbiased lipidomic analysis from indicated brain areas (control group subjects only). C) Heatmap shows Z scored relative lipid class abundance (control group) per brain regions. D) PCA plot of unbiased lipidomic analysis of AD (purple) and control (green) brain tissue samples from frontal cortex (FC) gray matter. E) Heatmap depicting changes (log2fold AD subject vs average control group) at the lipid class level for each individual AD subject and each brain area. AD patient samples are ordered 1-20 from left to right in each brain area (see methods for metadata) F) Average log2fold change of lipid classes in all AD brain samples compared to control samples per brain area. G) Changes in levels of CE, DG and TG (neutral) lipid species in control versus AD group. Mann-Whitney U test with Benjamini-Hochberg correction *=P<0.05. All lipid values in this figure are plotted as % of total lipids, raw concentration can be found in Supplementary figure 2.

### ApoE4 drives CE accumulation in human iPSC-derived astrocytes

CE accumulation drives pTau buildup in human neurons^7^ and alters microglial function after a myelin challenge^24,26^. While CE levels are highest in human astrocytes (Fig 1), the role of cholesterol esterification in these cells is unknown. Moreover, astrocytes express high levels of the AD risk gene APOE. A common variant in ApoE, ApoE4, is the major genetic risk factor for AD, and depending on ethnicity increases the risk for AD from 3-4 fold (heterozygosity) to 14-fold (homozygosity)^51,52^. AD pathology develops in virtually all ApoE4 homozygous carriers after the age of 65^53^. To map how ApoE4 affects the astrocytic lipidome, we differentiated two independent isogenic pairs of APOE3/3 and APOE4/4 iPSCs to astrocytes. We selected one isogenic pair from the iPSC Neurodegenerative Disease Initiative (parental line Kolf2.1J, APOE3/3, edited line APOE4/4 Kolf2.1J C156R hom3 male)^54,55^ and a second pair from the European Bank for Induced pluripotent Stem Cells (parental line BIONi037-A, APOE3/3, edited line BIONi037-A4, APOE4/4)^39^ (Fig 3A). Neither of these isogenic pairs has been characterized before by lipidomic and/or proteomic profiling. ApoE genotype of the iPSCs was confirmed by sanger sequencing on receipt and after differentiation to astrocytes, as well as after each experiment to verify sample identity (Sup Fig 3A). Successful differentiation to astrocytes was validated by astrocyte marker staining and expression of astrocyte markers by RNAseq^56^ (Fig 3B, Sup Fig 3B-C). ApoE4 astrocytes secreted less ApoE (Fig 3C), as has been reported in iPSC-derived astrocytes and CSF^5,8,10,57,58^. We performed lipidomics on two biological replicates (with 3 replicate samples per experiment) for both isogenic sets using our standardized iPSC-lipidomic pipeline (see Fig 3D-G and Sup Fig 3D-E for individual replicates, and Fig 3H and Sup Fig 3F for group level results). Strikingly, and consistently across experiments and lines, we observed a strong ApoE4-dependent increase in CE (individual species Fig 3D-G, Sup Fig 3D-E, and class level Fig 3H, Sup Fig 3F). Multiple TG species were also significantly increased, with near significance at the class level (Fig 3D-H). TGs containing saturated or monounsaturated fatty acids as well as highly polyunsaturated fatty acids (>5 double bonds) were most upregulated in our ApoE4 astrocytes (Fig 3I). Consistent with higher levels of storage lipids, lipid droplets were increased in ApoE4 astrocytes (Fig 3J). SM levels were significantly downregulated in ApoE4 astrocytes at both the species and class level, while LPE levels were only increased at the class level (Fig 3D-H, Sup Fig 3D-F). The (female) BIONi037 ApoE4 astrocytes also showed a strong and consistent increase in LacCER, HexCER and ceramide (CER) species, which was not found in the (male) Kolf2.1J ApoE4 astrocytes (Fig 3D-H, Sup Fig 3D-F). Of note, our study is not sufficiently powered to understand whether this is a haplotype, gender or line specific effect. We did not find evidence for increased saturation of phospholipids in our ApoE4 astrocytes, as is typical for activated astrocytes (Sup Fig 3G)^27^. Overall, our data indicates that ApoE4 strongly drives the accumulation of CEs, and to a lesser extent TGs and LPEs in human iPSC-derived astrocytes while decreasing SM levels. All lipidomics data of our ApoE4 and ApoE3 astrocytes are available on the Neurolipid Atlas.

**Figure 3.**
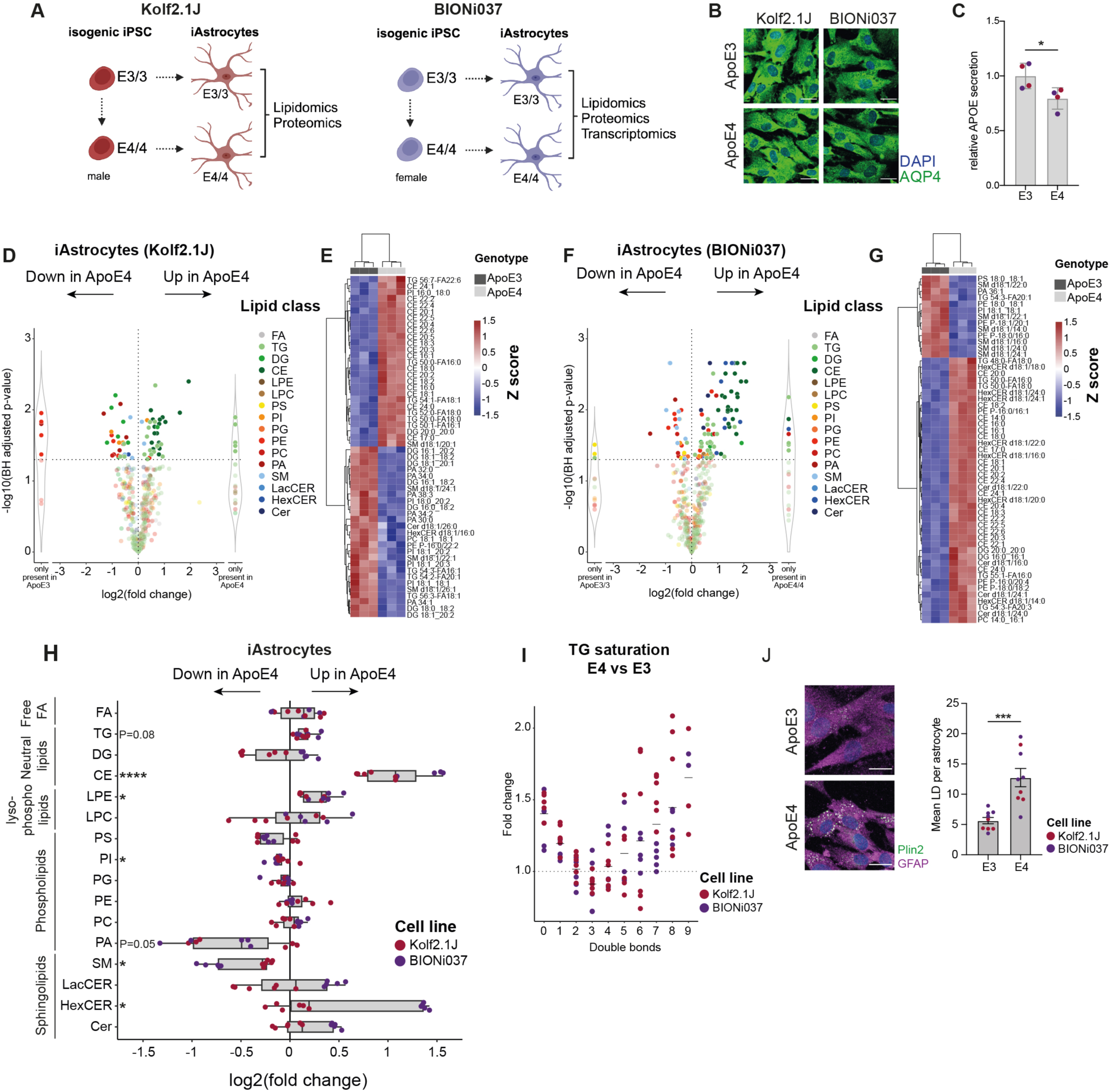
Lipidomic analysis of human isogenic APOE3/3 and APOE4/4 iPSC-derived astrocytes. A) Schematic overview of our multi-omics workflow with two independent isogenic iPSC-lines. B) Representative image of differentiated iAstrocytes from BIONi037 and Kolf2.1J background. Scale bar = 25mm C) Relative ApoE level secreted in the medium. n=4 wells from 2 independent lines. Mean + SD. *=P<0.05 t-test. D-G) Log2fold change of altered lipid species in BIONi037 (D) and Kolf2.1J (F) ApoE4 vs ApoE3 iAstrocytes. n=3 wells per genotype and heatmap of most differentiating lipid species between ApoE4 and ApoE3 iAstrocytes in BIONi037 (E) and Kolf2.1J lines (G) H) Summary data of changes in all detected lipid classes in ApoE4 vs ApoE3 iAstrocytes. n=6 samples per iPSC-line from 2 independent lipidomics experiments. Mann-Whitney U test with Benjamini-Hochberg correction *=P<0.05. I) Fold change in Triacylglycerides with indicated number of double bonds (unsaturation) in ApoE4 vs ApoE3 iAstrocytes. J) Representative image and quantification of the average lipid droplet number per astrocyte based on Plin2 staining. n=9 wells with E3 and E4 astrocytes (n=5 BIONi037, n=4 Kolf2.1J combined) from 2 independent experiments. Each datapoint represents a mean of >500 cells per well. Mean + sem. ***=P<0.0001 t-test.

### ApoE4 decreases interferon-dependent MHC class I antigen presentation and immunoproteasome pathways in human astrocytes

Our data for the first time indicate that ApoE4 increases CE levels in human astrocytes, but the functional consequence of this is not known. Therefore, we performed proteomic (both isogenic sets) and transcriptomic (only BIONi037 isogenic set) analysis on our ApoE4 versus ApoE3 astrocytes (Fig 3A, Fig 4A-C). Notably, the lipidomics in Figure 3, as well as the proteomics and transcriptomics were done on the same batch of astrocytes (see methods) to allow for multi-omic integration. We found 348 and 959 differentially expressed proteins (DEPs) in respectively Kolf2.1J and BIONi037 ApoE4 astrocytes (Fig 4A-C). ApoE was among these DEPs, showing downregulation in the ApoE4 astrocytes (Fig 4D). We focused our analysis on proteins that were either downregulated or upregulated in both ApoE4 lines (Figure 4E-F). Through overrepresentation analysis (ORA) we found that cell adhesion (e.g. NCAM1 interactions, integrin cell surface interactions) and extracellular matrix (ECM) related pathways (e.g. collagen chain trimerization, ECM proteoglycans) were upregulated by ApoE4 in both isogenic lines (Fig 4E). On the contrary, immune pathways (e.g. immunoregulatory interactions between a lymphoid and non-lymphoid cell, MHC I antigen presentation, interferon signaling) were downregulated (Fig 4F). This includes the term endosomal/vacuolar pathway which contained mainly MHC terms (Fig 4F). Strikingly, proteins in the MHC class I antigen presentation pathway were consistently downregulated as were immunoproteasome subunits, two pathways directly downstream of interferon signaling (Fig 4G). As a confirmatory read-out we stained against HLA class I heavy chain and confirmed downregulation of MHC class I by western blot (Fig 4H), immunohistochemistry (Fig 4I) and flow cytometry (Fig 4J). To be able to compare our results to previous transcriptomic studies with different ApoE4 iPSC-derived astrocytes^8,10^, we also performed transcriptomic analysis on our BIONi037 isogenic set (Sup Fig 4C-E). Through unbiased gene set enrichment analysis, the MHC class I antigen presentation pathway was shown to be downregulated by ApoE4 also at the transcriptome level (Fig 4K, Sup Fig 4E). Our transcriptomic analysis additionally showed downregulation of interleukin and interferon immune signaling pathways, the complement cascade and ER phagosome transport, while translation related terms were upregulated (Fig 4K, Sup Fig 4E). When looking at ApoE4 dependent gene expression changes in the complete Interferon reactome pathway (Sup Fig 4G) we found that similar to the proteomic results, specifically all genes in the MHC class I pathway and immune specific subunits of the proteasome, were downregulated by ApoE4 in our isogenic sets (Fig 4L) as well as in all isogenic sets and case-control sets from Lin *et al*. 2018 and TCW *et al*. 2022^8,10^ (Fig 4G,L). Changes in other pathways such as the complement cascade and translation initiation were observed, but the direction of change was highly variable across lines (Fig 4K, see discussion). Overall, these data show that ApoE4 decreases interferon signaling dependent pathways such as MHC class I antigen presentation and the immunoproteasome in human iPSC-derived astrocytes.

**Figure 4.**
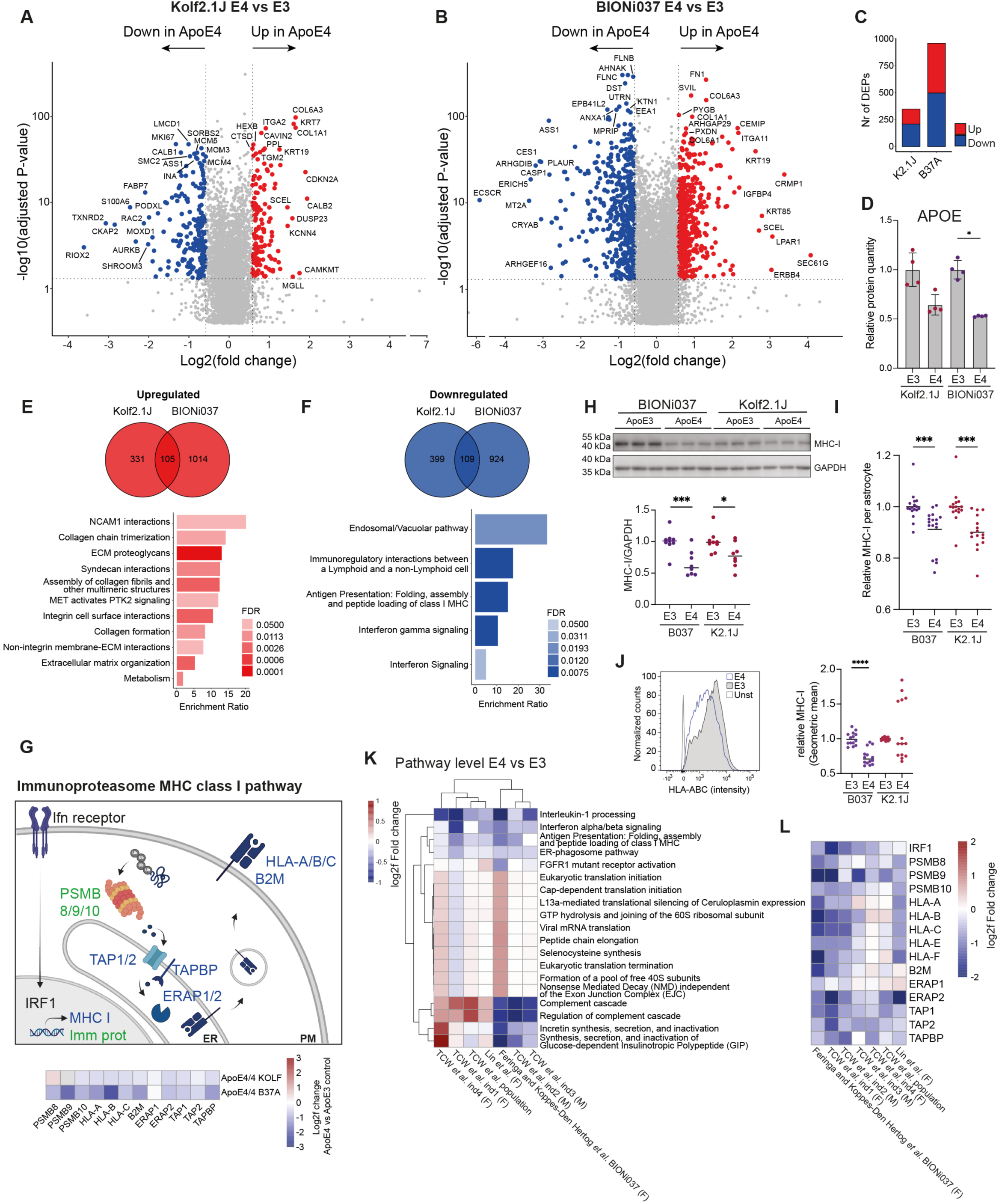
Proteomic and transcriptomic analysis of human isogenic APOE3/3 and APOE4/4 iPSC-derived astrocytes. A-B) Log2fold changes in protein levels in ApoE4 vs ApoE3 iAstrocytes from Kolf2.1J (A) and BIONI037 (B). Top ten proteins with highest log2fold change and top ten proteins with most significant P-value are labeled. n=4 wells per genotype. C) Number of DEPs (fold change > 1.5 & FDR <0.05) detected in ApoE4 vs ApoE3 iAstrocytes of Kolf2.1J or BIONi037 isogenic sets. D) Relative ApoE protein levels in ApoE3 and ApoE4 iAstrocytes (from proteomic analysis) from BIONi037 and Kolf2.1J background. Mean + SD *P<0.05 Mann-Whitney U test. E-F) Venn diagrams depicting the number of DEPs significantly upregulated (E) or downregulated (F) >1.25 fold times (>0.3 log2fold) in Kolf2.1J, BIONI037 or both ApoE4 iAstrocytes. A reactome overrepresentation analysis was performed on the 105 common upregulated (E) or 109 common downregulated (F) proteins and the enrichment ratio was plotted for all significant pathways (FDR <0.05). G) Schematic overview of interferon-dependent regulation of MHC class I antigen presentation (in blue) and immunoproteasome (in green) pathways. Heatmap indicated log2fold change of indicated proteins in ApoE4 vs ApoE3 iAstrocytes. PM = plasma membrane, ER = endoplasmic reticulum. H) Representative western blot and quantification of MHC-I levels (anti-HLA Class I Heavy Chain) in ApoE4 van ApoE3 iAstrocytes. BIONi037 (B037) and Kolf2.1J (K2.1J). n=8 wells from 3 independent experiments per line. Mean ***P<0.001 Mann-Whitney U test. I) Quantification of intracellular MHC-I levels as measured by immune fluorescence microscopy (stained for HLA-A,B,C). n=17 (B037) n=18 (K2.1J) wells from 6 independent experiments per line. Mean ***P<0.001 Mann-Whitney U test. J) Representative histogram (BIONi037) and quantification of plasma membrane MHC-I levels (stained for anti-HLA Class I Heavy Chain) by flowcytometry. Unst = unstained control. n=14 wells from 6 independent experiments per line. Mean ***P<0.001 Mann-Whitney U test. K) Comparison of significant reactome pathways (by gene-set enrichment analysis) from our transcriptomic analysis of ApoE4 vs ApoE3 (BIONi037) astrocytes with previously published datasets. Shown is the average log2fold change of all genes in the indicated pathway. TCW *et al.* ind1-4 (four different isogenic sets) and population (ctrl vs ApoE4 subjects) represent iPSC-derived astrocytes from ^10^, Lin *et al.* represents one isogenic set of ApoE4 vs ApoE3 iPSC-derived astrocytes from ^8^. (F)=Female (M)=Male L) Heatmap shows the log2fold change in individual genes in the MHC I and immunoproteasome pathway across indicated studies, including our data here. (F)=Female (M)=Male

### Reactive human astrocytes decrease CE levels, increase MHC I antigen presentation and immunoproteasome pathways

The reduction in interferon-dependent pathways is striking, as ApoE4 is thought to enhance, not decrease, immune signaling^5,10,59–62^. Yet our data clearly demonstrate a consistent reduction in the expression of proteins in these pathways including all (5) class I leukocyte antigens (HLA; HLA-A, HLA-B, HLA-C, HLA-E, HLA-F) and all specific subunits of the immunoproteasome (PSMB8/9/10) (Fig 4G,L, Fig 5K). To better understand our lipidomic and proteomic findings in the context of astrocyte immune function, we also performed multi-omic analysis of activated (TNF/Il-1α/C1q) iPSC-derived astrocytes (Kolf2.1J and BIONi037-A, Fig 5A). In contrast to the ApoE4 astrocytes, CEs and TGs were strongly downregulated in reactive astrocytes at the species (Fig 5B, Sup Fig 5A-E) and class level (Fig 5C,J, Sup Fig 5B-D), whereas HexCER were increased in reactive astrocytes as well as in the BIONi037 ApoE4 astrocytes (Fig 5C,J). As expected, phospholipid saturation was increased in activated astrocytes (Fig 5D, Sup Fig 5F)^27^, but not in ApoE4 astrocytes (Sup Fig 3G). Comparison of these lipid profiles indicates that the lipidome of ApoE4 and reactive astrocytes is distinct, especially with respect to stored lipids (CEs and TGs). These storage lipids increase in ApoE4 astrocytes but decrease in reactive astrocytes. Similar as above for ApoE4, we performed proteomics on the activated astrocytes to complement our lipidomics insights (Fig 5E-I). We found 431 and 469 differentially expressed proteins (DEPs) in respectively Kolf2.1J and BIONi037 activated astrocytes (Fig 5E-G). We focused our analysis on proteins that were upregulated or downregulated by reactivity in both lines (Figure 5H-I, Sup Fig 6A). Overrepresentation analysis did not identify any significant downregulated pathways (Sup Fig 6A). However, the topmost upregulated pathways (Fig 5H) were endosome/vacuolar pathway, immunoregulatory interactions between a lymphoid and non-lymphoid cell, MHC I antigen presentation and interferon signaling. Pathways that were all downregulated in ApoE4 astrocytes (Fig 4F). Beyond these terms, analysis showed that virtually all immune upregulated proteins in reactive astrocytes were down in the ApoE4 astrocytes (Fig 5K, Sup Fig 6B-C). By flow cytometry we confirmed that MHC class I levels (HLA class I heavy chains) were indeed increased in activated astrocytes (Fig 5L). Overall, our results indicate that ApoE4 and reactive astrocytes have opposing lipidomic and proteomic phenotypes. CEs and TGs are up in ApoE4 astrocytes, but down in reactive astrocytes, whereas interferon signaling-dependent pathways, the immunoproteasome and MHC class I are down in ApoE4 astrocytes but upregulated in reactive astrocytes (Fig 5M).

**Figure 5.**
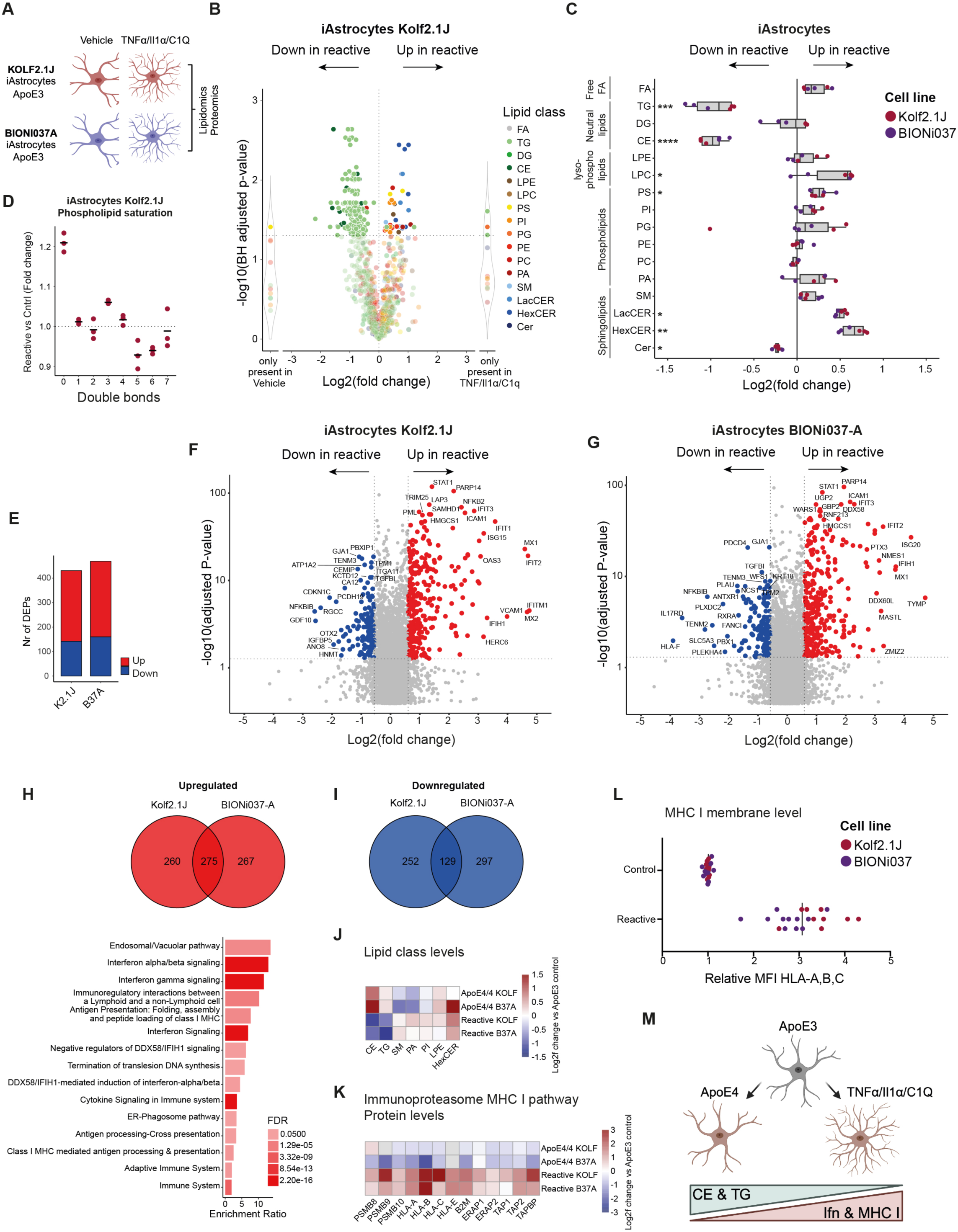
Lipidomics and proteomic analysis of reactive human iPSC-derived astrocytes. A) Schematic overview of experimental design, a cocktail of TNF/Il-1α/C1q was added for 24h hours to make astrocytes reactive. B) Log2fold change of altered individual lipid species in reactive vs control iAstrocytes (Kolf2.1J ApoE3). n=3 wells per condition. C) Fold change of all phospholipid species with indicated number of double bonds (unsaturation) in reactive vs control iAstrocytes (Kolf2.1J ApoE3). D) Summary data of changes in all detected lipid classes in reactive vs control iAstrocytes. n=3 wells per line. Mann-Whitney U test with Benjamini-Hochberg correction *=P<0.05. E) Number of DEPs (fold change > 1.5 & FDR <0.05) in reactive vs control iAstrocytes for indicated lines F-G) Log2fold changes in protein levels of reactive vs control iAstrocytes for Kolf2.1J (F) and BIONi037 (G). Top ten proteins with highest log2fold change and top ten proteins with highest P-value are labeled. n=4 wells per genotype. H) Venn diagram depicting the number of proteins that were significantly upregulated (H) or downregulated (I) >1.25 fold (>0.3 log2fold) in reactive Kolf2.1J, BIONi037 and both iAstrocytes. A reactome overrepresentation analysis was performed on the 275 common upregulated or 129 common downregulated proteins. No significantly enriched downregulatd pathways were observed, the enrichment ratios for all significantly (FDR<0.05) upregulated pathways are plotted in H. J) Heatmap depicting the log2fold change of indicated lipid classes (changed in ApoE4 iAstrocytes with P<0.1) in ApoE4 or reactive astrocytes vs ApoE3 control iAstrocytes. K) Heatmap depicting the log2fold change of indicated proteins from the MHC class I and immunoproteasome pathway in ApoE4 or reactive astrocytes vs ApoE3 control iAstrocytes. (Based on proteomics data) L) Relative membrane MHC-I levels (stained for anti-HLA Class I Heavy Chain) by flowcytometry in reactive vs control iAstrocytes. n=11 wells BIONi37 from 5 independent experiments and n=9 wells Kolf2.1J from 4 independent experiments. ****p<0.0001 unpaired t-test. M) Schematic representation of opposing lipidomic and proteomic phenotypes in ApoE4 and reactive iAstrocytes.

**Figure 6.**
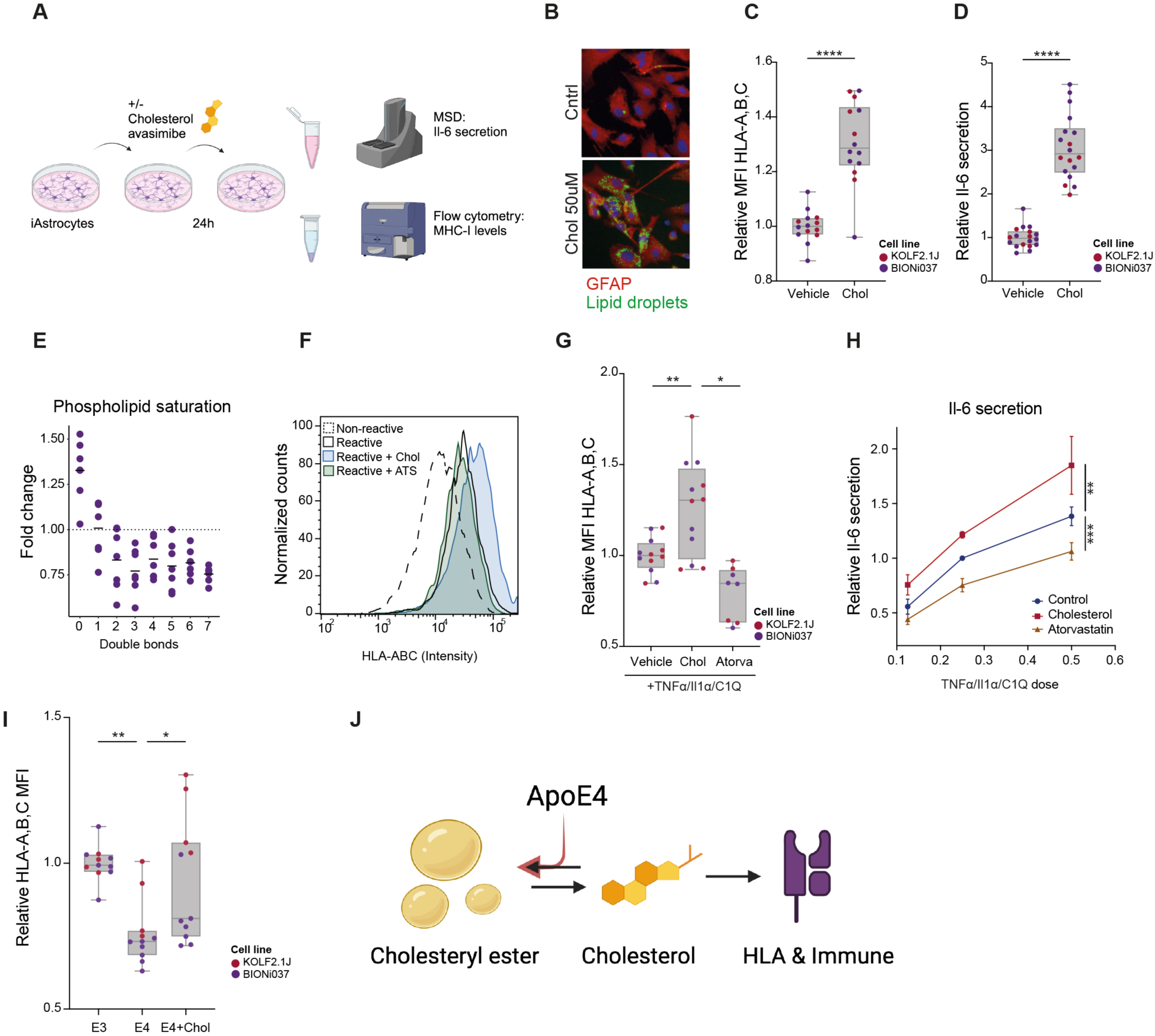
Cholesterol regulates activation of human astrocytes. A) Schematic representation of the experimental design. B) Lipid droplet staining in iAstrocytes following 24h treatment with cholesterol. C) Normalized membrane MHC-I levels (stained for anti-HLA Class I Heavy Chain) in vehicle versus cholesterol treated ApoE3 iAstrocytes determined by flow cytometry. n=6 (K2.1J) and n=8 (B037) from 4 independent experiments. ****P<0.0001 Unpaired t-test. D) Normalized Il-6 secretion in vehicle versus cholesterol treated ApoE3 iAstrocytes n=6 (K2.1J) and n=12 (B037) from 4 independent experiments. ****P<0.0001 Unpaired t-test. E) Fold change of phospholipid species with indicated number of double bonds (unsaturation) in cholesterol treated vs control iAstrocytes (BIONi037 ApoE3). n=6 wells from 3 independent experiments. F-G) Representative histogram and quantification (G) of normalized MHC-I membrane levels determined by flow cytometry (stained for anti-HLA Class I Heavy Chain) in response to indicated treatment conditions in iAstrocytes. n=6 (K2.1J) and n=6 (B037) wells from 3 independent experiments per line. *P<0.05 One-way ANOVA with Dunnett’s multiple comparison correction. H) Secreted Il-6 levels in medium of ApoE3 iAstrocytes that were pre-treated with vehicle, exogenous cholesterol (10mM) or atorvastatin (0.5mM for one hour and then treated for 24 hours with increasing doses of TNF/Il-1α/C1q (in presence of vehicle, atorvastatin or exogenous cholesterol). n=5 biological replicates (n=2 Kolf2.1J and n=3 BIONi037). **p<0.01 intercept difference by linear regression model. Relative Il-6 levels with vehicle 0.25 times cocktail dose set at 1. I) Relative changes in membrane MHC-I levels determined by flow cytometry (stained for anti-HLA Class I Heavy Chain) in ApoE3 or ApoE4 iAstrocytes treated with cholesterol. n=6 (K2.1J) and n=6 (B037) wells from 3 independent experiments. BIONi037 (B037) and Kolf2.1J (K2.1J). J) ApoE4 decreases HLA expression and immune function in human glia by increased cholesterol storage in cholesteryl esters.

### Cholesterol metabolism regulates MHC class I presentation and immune activation in human astrocytes, which is impaired by ApoE4

Based on the reduction of CEs in reactive astrocytes, but increase in ApoE4 astrocytes, we hypothesized that changes in cholesterol metabolism might directly contribute to immune phenotypes. We also found that a specific cluster of cholesterol synthetic genes was upregulated in reactive astrocytes (Sup Fig 7A). To test whether cholesterol regulates immune activation in human astrocytes, we treated Kolf2.1J and BIONi037-A control (ApoE3) astrocytes with cholesterol (Fig 6A-D). Both MHC class I presentation and IL-6 secretion were significantly increased after cholesterol treatment (Fig 6C-D). CEs are generated through conjugation of free cholesterol to a fatty acid by acyl coenzyme A-cholesterol acyltransferases (ACATs). Combined addition of cholesterol with the ACAT inhibitor avasimibe further increased IL-6 secretion, indicating that CE formation buffered the immune activation by cholesterol treatment (Sup Fig 7B). Moreover, the addition of cholesterol to astrocytes is sufficient to increase saturated phospholipid levels (Fig 6E) and the LPC lipid class level (Sup Fig 7C-D) typical for activated astrocytes (Fig 5C-D). Exogenous cholesterol also potentiated immune activation (as measured by MHC class I levels and IL-6 secretion) of astrocytes treated with TNF/Il-1α/C1q (Fig 5F-H). Conversely, pretreatment with atorvastatin (to reduce cholesterol levels) inhibited MHC class I upregulation and IL-6 secretion upon astrocyte activation by this cytokine cocktail (Fig 5F-H). Finally, we found that the addition of free cholesterol rescued MHC I expression in the ApoE4 astrocytes (Fig 6I). Overall, our data indicate that cholesterol is a major regulator of MHC class I antigen presentation and immune activation in human astrocytes. Furthermore, our data indicate that ApoE4 impairs astrocyte immune activation through increased esterification and storage of cholesterol in CEs (Fig 6J).

**Figure 7.**
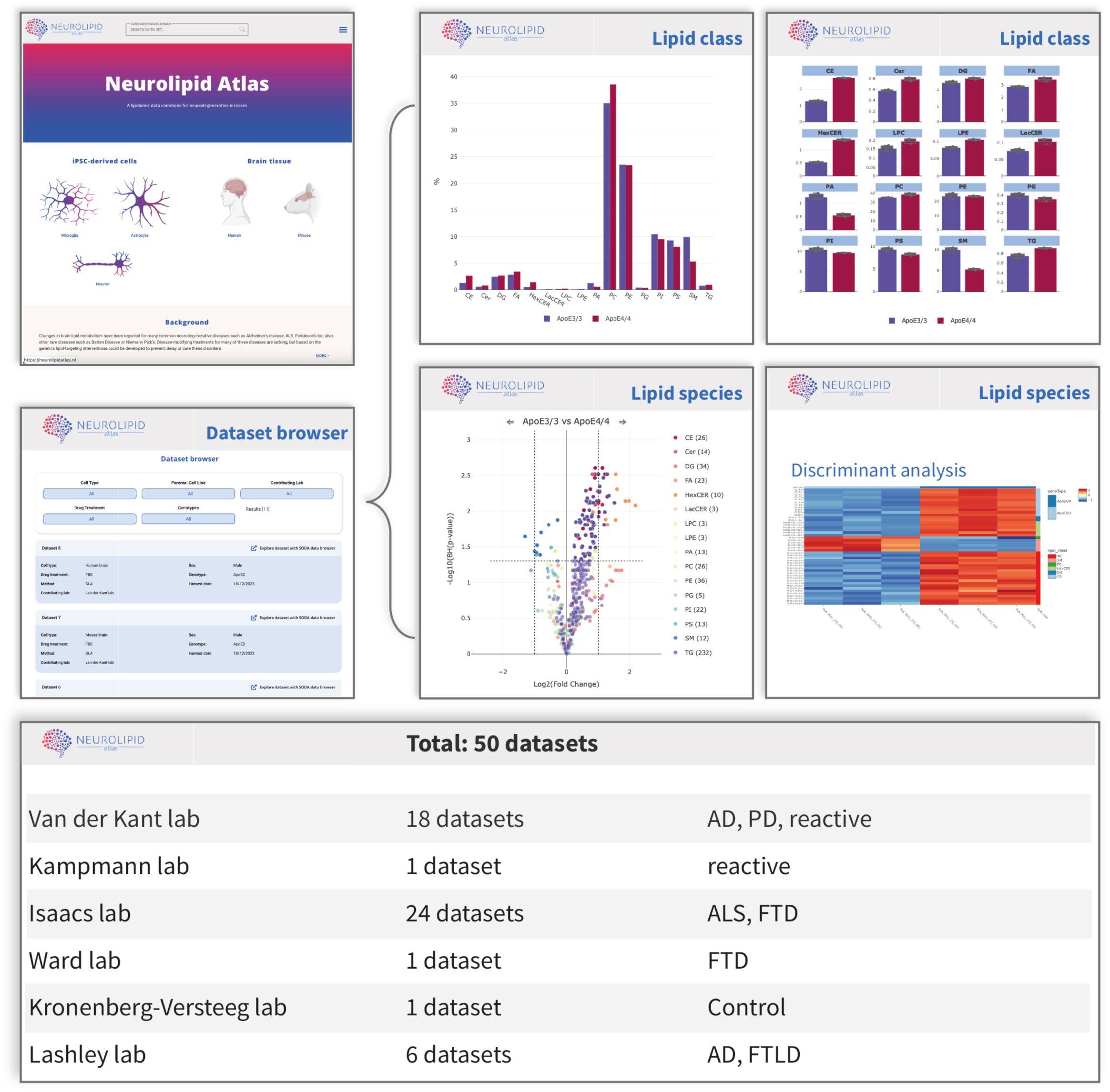
The neurolipid Atlas. Overview of the Neurolipid Atlas data commons (https://neurolipidatlas.com) to explore all lipidomics datasets from this study. Representative images of the start page, data browser as well as examples of bargraphs, volcano plot or heatmap for visualization of changes in lipid class or species levels between selected conditions. In addition, a summary list of currently available datasets is shown.

### The Neurolipid Atlas: an open-access lipidomics data commons for neurodegenerative diseases

We have generated an online lipidomics browser (Neurolipid Atlas, found at www.neurolipidatlas.com) that makes it easy to explore lipidomics data without prior bioinformatics knowledge (Fig 7). We have uploaded our data presented above, as well as (un)published lipidomics data generated together with a large group of collaborators, totaling a current number of 50 datasets over four neurodegenerative diseases and multiple treatment conditions (Sup Table S1). New datasets from our lab, and our collaborators, will be uploaded to the database in a continuous manner, and we invite other labs to contribute their lipidomics data to the Neurolipid Atlas. The Neurolipid Atlas allows for the download of all raw data and metadata, as well as the in-browser analysis, which includes quality control, blank filtering, normalization and the generation and customization of figures. Datasets can be searched for by name, cell type, genotype, treatment type, parental line or contributing lab. Users can explore and visualize changes at the lipid class level (bar graphs) or at the species level (volcano plots, heatmaps, principal component analysis, fatty acid analysis) and interact with the data by hovering over different lipid species. All figures, as well as their source data, can be downloaded.

## Discussion

Lipid metabolism is affected and implicated in various neurodegenerative diseases^2,3,14–19,21,22,24,26,4,6–12^. Here we generated a novel resource, the Neurolipid Atlas, to facilitate insights into lipid changes associated with different neurodegenerative diseases in a disease- and cell type specific manner. As proof of concept, we show that iPSC-derived neurons, astrocytes and microglia have distinct lipotypes that mimic their *in vivo* lipidomes. Furthermore, by comparative analysis on lipidomes of AD brain and iPSC-derived isogenic ApoE3, ApoE4 and reactive astrocytes from multiple donors we show that CE accumulation is a hallmark of AD and ApoE4 genotype, and that increased cholesterol esterification in ApoE4 astrocytes represses their immune function, specifically interferon pathways, MHC class I antigen presentation and immunoproteasome pathways.

### Cholesterol ester accumulation as a hallmark of Alzheimer’s disease

Our findings here further consolidate the notion that CE accumulation is a major pathogenic hallmark of AD^4,47,48^. CEs in CSF have also been shown to correlate with progression from mild cognitive impairment to AD^63^. CE accumulation in neurons drives pTau accumulation and Aβ overproduction^7,64–66^. Accumulation of CEs has also been detected in microglia upon TREM2 or ApoE loss^26^ and inhibition of CE formation improved Aβ clearance^67^. In ApoE4 oligodendrocytes CE accumulation led to perturbed neuronal myelination^68^. We now show for the first time that enhanced CE accumulation in astrocytes represses their immune function and the MHC class I pathway. An important next step would be to determine which cell type(s) in the AD brain accumulate these CEs in human post-mortem material. Unfortunately, lipidomics techniques do not currently have enough resolution to identify CE at the single cell level. Our data show CEs accumulate in both white and gray matter of the frontal cortex, which indicates that cells other than oligodendrocytes^68^ also contribute to this phenotype. Secondary neurodegenerative processes (protein aggregation, neuronal death, demyelination) and CE accumulation in microglia might be a possible explanation for the observed CE accumulation in AD subjects^24–26^. However, our data also indicates an increasing trend in CE levels in the cerebellum of AD patients which is relatively spared from late-stage pathology in AD. Also, our finding that the AD genetic risk factor ApoE4 strongly drives CE accumulation in astrocytes in the absence of pathology, indicates that CE accumulation might not merely be a downstream effect of neurodegeneration, but rather is directly downstream of AD risk genes. Increases in TG levels have been reported in the aging mouse brain^23^, and we and others^9^ also observed an increase in TGs in ApoE4 astrocytes as well as a trend towards increased TG levels in the AD brain (Fig 2). We confirm previous reports that ApoE4 increases levels of polyunsaturated triglycerides^9,69^, but also find an increase in monounsaturated TG levels (Fig 3I). Importantly, ApoE4 astrocytes do not show the increase in saturated phospholipids typical for reactive astrocytes^27^ (Sup Fig 3G). Overall, our data show that ApoE4 and AD present a unique lipotype that is primarily characterized by strong CE accumulation, as well as changes in TGs and SM.

Lowering cholesterol esterification has been shown to have beneficial effects in many AD models^64,67,70–73^ including human iPSC-derived neurons^7^, oligodendrocytes^68^, microglia^26^ and now also astrocytes (Fig 6). However, ACAT inhibition (by Pactimibe^74,75^ or Avasimibe^76^) has failed in clinical trials for the treatment of cardiovascular disease for lack of effect^75,76^ or increased cardiovascular events^74^. No FDA-approved ACAT-inhibitors exist. Though, our work here on CE and the body of evidence supporting beneficial effects of ACAT inhibition on AD pathogenesis in a number of brain cell types, encourages future consideration of ACAT inhibitors as possible treatment strategy for AD, especially for ApoE4 carriers.

### iPSC modeling of ApoE4 effects in astrocytes

As the strongest genetic risk factor for AD (ApoE) is highly expressed in astrocytes, there is an urgent need to understand how the AD risk variant ApoE4 affects astrocytes. Here we provide the first characterization of an isogenic APOE3/3 and APOE4/4 pair of iPSC-derived astrocytes from the INDI line Kolf2.1J which we hope will serve as a reference to the field. We also provide the first full characterization of a second isogenic pair of iPSC-derived astrocytes (BIONi037, EBISC). Our study highlights that there is significant variability between parental lines. For example, LacCER, HexCER and ceramide were strongly increased in the female BIONi037 ApoE4 line, but not in the male Kolf2.1J ApoE4 line. Similarly, our proteomics analysis (Sup Fig 4A-B) revealed several differences between ApoE4 effects in the different parental lines with stronger overall effects in the female BIONi037 ApoE4 line. This could indicate a gender, haplotype or clone specific effect. It is important to stress that to date iPSC-studies aimed at ApoE4, even with isogenic lines, have not been powered to address (genetic) context-specific differences and more studies are needed. For example, when comparing our transcriptomics data with two previously published transcriptomic datasets on ApoE4 astrocytes we find that pathways such as the complement system and RNA processes are up in some lines and down in others. In strong contrast, interferon signaling, MHC class I antigen presentation and ER-phagosome pathways were down in ApoE4 lines from all these studies (Fig 4) indicating a robust and likely context independent effect of ApoE4 on the suppression of immune pathways. In addition, our proteomics data showed upregulation of extracellular matrix related pathways in ApoE4 astrocytes, which was also recently described in ApoE4 astrocytes by TCW *et al.* 2022^10^. Interestingly, another recent study found rare coding variants in extracellular matrix genes to protect against AD development in APOE4 carriers^77^.

### ApoE4 immune suppression

Astrogliosis is a major feature of end-stage AD^78–81^. We were therefore very surprised to find major immune pathways such as interferon signaling, the immunoproteasome and MHC class I antigen presentation to be downregulated in ApoE4 astrocytes. To place this in context we also generated the first lipidomic and proteomic analysis of (TNF/Il-1α/C1q) activated human iPSC-derived astrocytes. We confirmed that astrocyte activation (as in mouse^27^) also induces phospholipid saturation in human iPSC-derived astrocytes. Strikingly, the topmost upregulated proteomic pathway in our reactive astrocytes were interferon pathways including MHC class I. Also, CE and TG were down in reactive astrocytes, but up in ApoE4 astrocytes (Figure 5J). The results provide strong evidence that ApoE4 intrinsically inhibits, rather than activates, astrocyte immune function. These results fit with recent reports in AD mice showing decreased immune function of ApoE4 microglia, including reduced antigen presentation^82,83^. Interestingly, human stem cell-derived microglia xenotransplanted into *APP*^NL-G-F^ mice were shown to transition into a human specific HLA expressing state and ApoE4 selectively reduced the proportion of cells acquiring this HLA phenotype^84^. In line with this, an AD protective variant in PLCg2 (PLCg2 P522R) has recently been shown to reduce CE accumulation in iPSC-derived microglia^24^ while increasing microglial MHC I levels and providing benefit through increased recruitment of T-cells^85^. Based on these data, the presence of a similar ApoE4-cholesterol-immune axis in microglia, as we identified here for astrocytes, is likely but needs to be confirmed. However overall, these findings (including ours in human brain cells) indicate that ApoE4 intrinsically limits immune activation, rather than inducing immune activation. This could indicate that immune activation (of astrocytes and microglia) is actually needed to stave-off AD, and that restoration (or activation) of glial activity in ApoE4 carriers might prevent AD pathogenesis. With current technologies the hypothesis that ApoE4 suppresses glial immune activation before AD onset is difficult to validate in human subjects as post-mortem material normally reflects late disease states. It would therefore be highly relevant to evaluate e.g. immunoproteasome levels, MHC class I expression and lipid levels in healthy ApoE4-carriers early in life, for example trough tissue obtained from normal-pressure hydrocephalus biopsies^86^. The exact pathway connecting cholesterol levels to interferon pathways, the immunoproteasome and MHC class I antigen presentation also needs more study, but may involve direct interaction between cholesterol and interferon signaling at lipid rafts^87,88^. However, our study is the first to show that the effect of ApoE4 on glial immune function is mediated by ApoE4-induced changes in glial lipid metabolism and storage.

Overall, our data highlight the important role of lipid- and particularly cholesterol-metabolism in AD. We created a novel tool (the Neurolipid Atlas) as a resource of lipidomic datasets for different cell types, mutations, neurodegenerative diseases and model organisms. As a proof of concept, we show that iPSC-derived neurons, astrocytes and microglia have distinct lipidomes that recapitulate *in vivo* lipotypes. Our data solidifies the link between AD and cholesterol, further establishing CE accumulation as a hallmark of AD. Finally, we show for the first time that cholesterol regulates astrocytic immune function, which is impaired by the genetic AD risk variant ApoE4.

## Methods

### iPSC culture

Isogenic Kolf2.1J (APOE3/3) and Kolf2.1J C156R Hom3 (APOE4/4) human iPSCs were a kind gift from INDi (Donor 57y male). Isogenic BIONi037-A (APOE3/3) and BIONi037-A4 (APOE4/4) human iPSC lines were obtained via EBISC (Donor 77y female). iPSCs were cultured in 6-well plates precoated with 120-180 µg/mL Geltrex (Fisher Scientific, A1413302) in Gibco™ Essential 8™ medium (E8; Fisher Scientific, 15190617) + 0.1% Pen/Strep (P/S; Fisher Scientific, 11548876), with daily full medium refreshments. iPSC colonies grown till 90% confluency were dissociated using 1 mM EDTA (Invitrogen, 15575-038) in 1X PBS (VWR, 392-0434) and replated in Essential E8 medium supplemented with 5 μM ROCK Inhibitor (RI; Tebu Bio, Y-27632). Genomic integrity of iPSC lines was periodically tested based on SNP arrays. In addition, cell cultures were regularly tested for mycoplasma contamination.

### Quality control cells

DNA from cell cultures was isolated using ReliaPrep gDNA Tissue Miniprep System (Promega, A2052). Samples were processed by the Global Screening Array (GSA) Consortium Project at Erasmus MC Rotterdam, The Netherlands on the Illumina GSA beadchip GSA MD v1. SNP data was processed and annotated with Illumina GenomeStudio software (Illumina, San Diego, CA). iPsychCNV package was used for copy number variant (CNV) calling, which integrates B allele frequency distribution and Log R ratio to reduce false positive detection (Bertalan, 2017). CNVs larger than 25 kB and containing more than 100 SNPs were flagged and compared against gene lists associated with brain development and synapse GO terms. In addition, DNA from iPSC-derived Astrocytes (iAstrocytes) in each experiment was isolated to confirm the APOE genotype.

### iPSC differentiation to neurons

NGN2 transcription-based iPSC differentiation to neurons was based on^40^. iPSCs were infected in suspension (in E8 + RI) with ultra-high titer lentiviral particles provided by ALSTEM, encoding pTet-O-Ngn2-puro (Addgene #52047) and FUΔGW-rtTa (Addgene #19780). To start neuronal induction 100K cells/cm2 infected iPSCs were plated in N2-supplemented medium (DMEM/F-12 + GlutaMAX (Thermo Fisher, 31331093), 3g/L D-glucose (Thermo Fisher, A2494001), 1% N2 supplement-B (Stemcell Technologies, 07156) and 0.1% P/S) supplemented with 5 µM RI, 2 μg/ml doxycycline hyclate (Sigma Aldrich, D9891) and dual SMAD inhibitors (100 nM LDN193189 (Stemgent, 04-0074), 10 μM SB431542 (Tebu-Bio, T1726), 2 μM XAV939 (Sigma-Aldrich, X3004)). On day 2, 100% of the medium was refreshed (including all day 1 supplements except RI), and 3 µg/mL puromycin was added (Cayman Chemical, 13884-25). On day 3, 100% medium was exchanged for N2-supplemented medium with doxycycline hyclate, puromycin and 10 µM 5-fluooro-2’-deoxyuridine (FUdR; Sigma-Aldrich, F0503). 6-well plates were coated with 20 µg/mL poly-L-Ornithin (PLO; Sigma-aldrich, P3655) overnight at room temperature (RT) followed by 3 wash steps with PBS on day 4. PLO-coated wells were subsequently coated with 5 µg/mL laminin (lam; Bio-techne, 3400-010-02) for 2-4h at 37°C. iPSC-derived neurons (iNeurons) were washed with 1X PBS before dissociation with accutase (Merck, SCR005) for 5 min at 37°C. iNeurons were collected in DMEM (VWR, 392-0415P) and pelleted by 5 min spin at 180g. iNeurons were resuspended and plated at 600K/well in PLO-lam coated 6-well plates in Neurobasal medium (NBM; Fisher Scientific, 11570556), supplemented with 200 mM GlutaMAX (Thermo Fisher, 35050038), 3 g/L D-glucose (Thermo Fisher, A2494001), 0.5% NEAA (Fisher Scientific, 11350912), 2% B27 (Fisher Scientific, 17504044), 0.1% P/S, 10 ng/mL BDNF (Stemcell Technologies, 17189321), 10 ng/mL CNTF (Peprotech, 450-13), 10 ng/mL GDNF (Stemcell Technologies, 78058.3). iNeurons were cultured at 37°C and 5% CO_2_ and medium was replaced with 50% fresh medium once a week.

### iPSC differentiation to astrocytes

iPSCs were differentiated to neuronal progenitor cells (NPCs) based on^89^. At day 1, iPSCs were plated at 100% density in 6-well plates in NMM medium (50% DMEM/F-12 + GlutaMAX (Thermo Fisher, 31331093), 50% NBM, 100mM GlutaMAX, 0.5% N2 supplement B, 1% B27, 0.5% ITS-A (Thermo Scientific, 51300044), 0,5% NEAA, 0.08% 2-mercaptoethanol (Fisher Scientific, 11528926) and 1% P/S supplemented with 10 µM SB431542 and 0.5 µM LDN193189. Complete medium was replaced daily for 7 days. At day 8, cells were expanded to PLO-lam coated 6 cm dishes. 1 mL EDTA per well was added after one PBS wash and cells were incubated at 37°C for 3-4 minutes. Cells were collected in clumps using a cell scraper and plated in 5 mL complete NMM medium supplemented with 5 μM RI. At day 9, medium was exchanged for plain NMM medium without inhibitors after one PBS wash. This medium was refreshed daily for 2 more days. At day 12, medium was exchanged for NMM medium supplemented with 10 ng/mL FGF (Peprotech, 100-18B). This medium was refreshed daily for 2 more days. On day 15, cells were incubated in accutase after one PBS wash for 5 min at 37°C and collected in NMM + 5 µM RI. After a 5 min spin at 1000 rpm, the pellet was resuspended in NMM supplemented with FGF and RI before plating the NPC cells (now P=1) in 2 PLO-lam coated 10 cm dishes. Medium was refreshed daily with NMM + FGF for the next 3 days. NPCs were maintained at high density and refreshed every 2-3 days. NPCs were plated for control stainings (Nestin/Pax6) at passage 4 to confirm NPC identity after which astrocyte differentiation was started based on Fong et al. (2018). One confluent 10 cm dish of NPCs was washed with 1X PBS before adding 9 mL NMM + FGF. Cells were collected in clumps by cell scraper and transferred at 3 mL/well to a non-coated 6-well plate. Plates were placed on an orbital shaker (90rpm) in a 37°C incubator. After 24h, when tiny neurospheres had formed 5 µM RI was added per well. 48h later, medium was changed back to NMM without FGF. One week after cell scraping of the NPCs the NMM medium was exchanged for astrocyte medium (AM; ScienCell, 1801) and afterwards medium was refreshed 3 times a week for the following 2 weeks. Neurospheres from 3 wells were collected and plated in 1 PLO-lam coated 10 cm dish. iAstrocytes differentiated from the neurospheres were passaged to uncoated 10 cm dishes using accutase and maintained in AM + 2% FBS (ScienCell, 1801/0010) until they were P4. iAstrocytes were plated for experiments when they were between P4 and P12.

### iPSC differentiation to microglia

iPSC-derived microglia (iMic) were generated following^35^ with small modifications. In brief, iPSCs were detached with Accutase (Gibco) and collected as single cell suspension. After centrifugation (5 minutes, 300 g, room temperature), 2.5 mil cells were plated into 24-well AggreWell800 plates (Stem Cell Technologies; pre-treated with Anti-Adherence rinsing solution) in 2 ml EB Induction Medium (mTeSR^+^ (Stem Cell Technologies) + 20 ng/ml SCF (R & D Systems) + 50 ng/ml BMP4 (Miltenyi) + 50 ng/ml VEGF (Miltenyi), supplemented with 10 μM Y27632 (Stem Cell Technologies) for the first 24 hours) per well to generate embryoid bodies (EB). To allow the formation of EBs, cells remained in AggreWell plates with daily 75% media changes for 5 days. After 5 days, EBs were harvested and equally distributed to two 6-well plates (Corning) in 2 ml of EB Differentiation medium (X-Vivo 15 (Lonza), 2 mM Glutamax (Gibco), 0.55 mM β-mercaptoethanol (Gibco), 100 U/ml/100 μg/ml Penicillin/Streptomycin (ThermoFisher Scientific), 25 ng/ml IL-3 (Miltenyi Biotec), 100 ng/ml M-CSF (Miltenyi Biotec)) per well. The EBs were kept in EB Differentiation Media at 37 °C and 5 % CO_2_ with full media changes every 7 days. After 2-3 weeks, non-adherent microglial precursor cells (pre-iMics) started to be released into the medium from EBs. Pre-iMics were harvested during regular medium changes by collecting the supernatant medium, strained through a 40 µm cell strainer (Greiner). Pre-iMics harvested in weeks 3-6 post-emergence were pooled and sustained in EB Differentiation Medium in T75 flasks (Corning) with weekly media changes. Once sufficient cell numbers had been collected, pre-iMics were plated at 15,000 cells/cm^2^ in T175 flasks (Sarstedt) in iMic Medium (50 % Advanced Neurobasal Medium (Gibco), 50 % Advanced DMEM-F12 (Gibco), 1 x B27 Supplement with Vitamin A (Gibco), 2 mM Glutamax (Gibco), 0.1 mM β-mercaptoethanol (Gibco), 100 ng/ml IL-34 (Miltenyi Biotec), 20 ng/ml M-CSF (Miltenyi Biotec)) and differentiated to iMics for 14 days. For each line 4 replicates were plated and processed in parallel. iMics were cultivated at 37 °C and 5 % CO_2_ with 3 full media changes per week. On Day 14, iMics were washed briefly with PBS, detached with Accutase for 6-7 minutes at 37 °C until cells detached upon tapping the flask. Cells were collected with Wash Buffer (Advanced DMEM-F12 (Gibco) + 0.1 % BSA Fraction V (Gibco)), centrifuged at 300 g, 5 min, RT before they were resuspended in PBS and counted using a hemocytometer (Neubauer Zählkammer Improved, Bard). Appropriate volumes containing 1 million cells were transferred to 1.5 ml microcentrifugation tubes (Eppendorf) and centrifuged at 400 g, 4 °C for 5 minutes. The supernatant was aspirated and cell pellets were frozen to −80 °C.

### Postmortem brain sample lipidomics

Lipidomic analysis was undertaken on human postmortem brain material including frontal cortex gray matter, frontal cortex white matter and cerebellum tissue from 13 control donors and 20 AD patient donors. Brain tissue was obtained from the Queen Square Brain Bank, UCL Queen Square Institute of Neurology. All donor information, including postmortem (PM) delay, age, sex, APOE genotype and pathological information is listed in table S1. Ethical approval for the study was obtained from the NHS research ethics committee (NEC) and in accordance with the human tissue authority’s (HTA’s) code of practice and standards under licence number 12198.

**Table S1.** Metadata human postmortem brain tissue.

| Sample nr | PM delay | AAO | AAD | Duration | Gender | Clinical Diag | Path Diag | Cleanedpat hdiag | Brain Weight | APOE | Braak Tau | Thal Phase | CERAD | ABC | CAA |
| --- | --- | --- | --- | --- | --- | --- | --- | --- | --- | --- | --- | --- | --- | --- | --- |
| 1 | 34:00:00 | 54 | 63 | 9 | M | Picks/FTD (bvFTD) | AD | AD | 1150 | 44 | 6 | 5 | 3 | A3B3C3 | 2 |
| 2 | 90:05:00 | 64 | 77 | 13 | M | AD (Amnesic) | AD | AD | 1264 | 44 | 6 | 5 | 3 | A3B3C3 | 3 |
| 3 | 60:25:00 | 59 | 76 | 17 | F | AD | AD | AD | 1191 | 44 | 6 | 5 | 3 | A3B3C3 | 2 |
| 4 | 70:00:00 | 55 | 68 | 13 | M | DLB | AD | AD | 1522 | 33 | 6 | 5 | 3 | A3B3C3 | 1 |
| 5 | 76:40:00 | 49 | 62 | 13 | F | AD (Amnesic) | AD | AD | 996 | 33 | 6 | 5 | 3 | A3B3C3 | 0 |
| 6 | 84:45:00 | 58 | 69 | 11 | M | PPA / AD | AD | AD | 1600 | 33 | 6 | 5 | 3 | A3B3C3 | 3 |
| 7 | 61:19:00 | 63 | 79 | 16 | M | AD | AD | AD | 1423 | 33 | 6 | 5 | 3 | A3B3C3 | 3 |
| 8 | 78:15:00 | 69 | 81 | 12 | M | AD | AD | AD | 1116 | 33 | 6 | 5 | 3 | A3B3C3 | 2 |
| 9 | 31:42:00 | 48 | 63 | 15 | M | AD | AD | AD | 1042 | 33 | 6 | 5 | 3 | A3B3C3 | 3 |
| 10 | 34:25:00 | 54 | 65 | 11 | M | AD | AD | AD | 1089 | 44 | 6 | 5 | 3 | A3B3C3 | 3 |
| 11 | 17:10:00 | 55 | 67 | 12 | F | PPA | AD | AD | 1009 | 33 | 6 | 5 | 3 | A3B3C3 | 3 |
| 12 | 14:50:00 | 62 | 72 | 10 | M | PD | AD | AD | 1180 | 44 | 6 | 5 | 3 | A3B3C3 | 2 |
| 13 | 66:00:00 | 51 | 60 | 9 | F | CBD | AD | AD | 1090 | 33 | 6 | 5 | 3 | A3B3C3 | 3 |
| 14 | 33:26:00 | 63 | 74 | 11 | M | AD | AD | AD | 1022 | 44 | 6 | 5 | 3 | A3B3C3 | 3 |
| 15 | 44:05:00 | 68 | 84 | 16 | F | VD | AD | AD | 1127 | 44 | 6 | 5 | 3 | A3B3C3 | 3 |
| 16 | 45:35:00 | 52 | 71 | 19 | M | AD | AD | AD | 1097 | 33 | 6 | 5 | 3 | A3B3C3 | 3 |
| 17 | 52:30:00 | 53 | 68 | 15 | F | Picks | AD | AD | 1103 | 44 | 6 | 5 | 3 | A3B3C3 | 1 |
| 18 | 40:10:00 | 49 | 69 | 20 | F | AD | AD | AD | 986 | 44 | 6 | 5 | 2 | A3B3C2 | 3 |
| 19 | 92:47:00 | 50 | 66 | 16 | F | Picks (PPA) (Amnesic) | AD | AD | 906 | 33 | 6 | 5 | 3 | A3B3C3 | 2 |
| 20 | 30:25:00 | 59 | 79 | 20 | F | AD | AD | AD | 961 | 33 | 6 | 5 | 3 | A3B3C3 | 3 |
| 21 | 80:00:00 |  | 64 |  | M | Control | Control | Control | 1695 | 33 | 0 | 1 | 0 | A1B0C0 | 0 |
| 22 | 47:00:00 |  | 73 |  | M | Control | Control | Control | 1291 | 33 | 4 | 1 | 0 | A0B2C0 | 0 |
| 23 | 171:00:00 |  | 69 |  | M | Control | Control/path aging | Control | 1435 | 33 | 1 | 3 | 1 | A2B1C1 | 0 |
| 24 | 26:46:00 |  | 70 |  | F | Dystonia? | Control | Control | 1200 | 23 | 1 | 0 | 0 | A0B1C0 | 0 |
| 25 | 79:00:00 |  | 76 |  | M | Control | Control | Control | 1366 | 34 | 2 | 1 | 0 | A1B1C0 | 0 |
| 26 | 24:00:00 |  | 73 |  | F | Control | Path Ageing | Control | 1214 | 34 | 2 | 2 | 2 | A1B1C2 | 1 |
| 27 | 76:10:00 |  | 71 |  | F | Control | Control | Control | 1214 | 33 | 3 | 2 | 1 | A1B1C0 | 2 |
| 28 | 47:05:00 |  | 89 |  | M | Control | Control | Control | 1356 | 33 | 2 | 3 | 1 | A2B1C1 | 1 |
| 29 | 29:30:00 |  | 78 |  | F | Control | Control age related changes | Control | 1225 | 22 | 2 | 0 | 0 | A0B2C0 | 0 |
| 30 | 88:50:00 |  | 79 |  | F | Control | Control | Control | 1288 | 33 | 1 | 2 | 1 | A2B1C1 | 0 |
| 31 | 45:05:00 |  | 68 |  | F | Control | Control | Control | 1330 | 23 | 0 | 0 | 0 | A0B0C0 | 0 |
| 32 | 54:20:00 |  | 73 |  | M | Control | Control | Control | 1498 | 24 | 3 | 4 | 2 | A3B2C2 | 3 |
| 33 | 3:30:00 |  | 79 |  | M | Control | Control | Control | 1355 | 33 | 2 | 0 | 0 | A0B1C0 | 0 |

Processing of postmortem samples for lipidomics was carried out as follows, after adding stainless steel beads and LC-MS grade water, brain samples were homogenized using a Next Advance bullet blender. From these homogenized samples, aliquots containing the equivalent of 5 mg of tissue were prepared as described below.

### Lipidomic analysis

Lipidomics analysis followed standardized, quantitative protocols (Ghorasaini et al., 2021, 2022). Briefly, 25 µL Lipidyzer internal standard mix containing 54 deuterated standards was added to the cell pellet and extraction followed a methyl tert-butyl ether-based protocol. After drying under a gentle stream of nitrogen, samples were dissolved in running buffer (methanol:dichloromethane 1:1, containing 10 mM ammonium acetate) and injected into the Lipidyzer platform, consisting of a SCIEX QTRAP 5500 mass spectrometer equipped with SelexION DMS interface and a Nexera X2 UHPLC-system. SLA software was used to process data files and report the lipid class and species concentration and composition values (Su et al., 2021). Lipidyzer data analysis was further accomplished using SODAlight as a built-in data browser for the Neurolipid atlas repository. Lipid species concentration datasets were imported and filtered, with individual species required to have a minimal intensity of two times the blank in at least 80% of all samples measured. If lipid species were absent, or below two times the blank in >20% of all samples, they were removed. An exception is made for lipid species that are uniquely present in one group, if within one of the experimental groups a lipid species is present in at least 60% of the samples (with a minimal intensity of two times the blank) the lipid species is reintroduced for the analysis. Because a data set can have several grouping variables (e.g. genotype, treatment, sample type) a new group variable is created by concatenating all grouping variables. This new group variable is used as the group variable for the blank filtering. No missing value imputation was done. The SLA control software, including all up to date dictionaries and isotope correction algorithms can be found here, https://github.com/syjgino/SLA. SODA-light is a development branch of iSODA [https://github.com/ndcn/soda-ndcn] and part of the Neurolipid Atlas.

### Neurolipidatlas

SODA-light was forked as a lipidomics-only instance of iSODA, a multi-omics data visualization and integration application developed on R 4.4.0. As such, SODA-light is designed for efficient data exploration, providing interactive plots with extensive flexibility in terms of input data, analytical processes, and visual customization. The code for SODA light is available at [https://github.com/CPM-Metabolomics-Lipidomics/soda-light]. SODA light version 0.2 was used for generation of all figures in this manuscript.

### Phospholipid and TG saturation analysis

To investigate differences in saturation of lipid classes between groups, the sum of the concentration of the lipid species with identical numbers of double bonds within the TGs or within all phospholipid classes was calculated. These summed values were normalized over total lipid concentration. Afterwards the fold change from each sample was calculated over the mean of the control samples.

### Experimental set-up for astrocyte multi-omics ApoE4 vs ApoE3

At day 1, iAstrocytes were plated at 17K cells/cm^2^ in AM + 2% FBS in uncoated 10 cm dishes. Both BIONi037-A (APOE3/3) and BIONi037-A4 (APOE4/4) or Kolf2.1J (APOE3/3) and Kolf2.1J C156R (hom3) (APOE4/4) were analyzed in 2 separate experiments, creating 4 datasets in total (BIONi037 set I & II and Kolf2.1J set I & II in the Neurolipid Atlas). At day 2, medium was replaced for AM without FBS after one PBS wash. After 24h, at day 3, iAstrocytes were collected by accutase dissociation after one PBS wash. iAstrocytes from each replicate dish were divided over 2 vials in which 500K −1 million iAstrocytes were collected for lipidomic and proteomic analysis. Cell pellets were stored at −80°C until shipment for further analysis. 4 (in case of proteomics) and 3 (in case of lipidomics) replicate samples per experiment were included. All lipidomic and proteomic data in figure 4 is from Kolf2.1J and BIONi037 iAstrocytes cultured without FBS. Proteomic analysis was done on Bioni037 set I and Kolf2.1J set II. Main figure 3 shows lipidomic analysis on Kolf2.1J set II and BIONi037 set II, while supplementary figure 3 shows lipidomic analysis of Kolf2.1J set I and BIONi037 set I.

In parallel this experimental set-up was performed in iAstrocytes that were cultured in AM + 2% FBS throughout the experiment. In that case iAstrocytes were collected at day 3 and divided over 3 vials for lipidomic, proteomic and transcriptomic analysis. The comparison of proteomic and transcriptomic analysis shown in Sup Fig 4C, as well as all transcriptomic data are from the BIONi037 iAstrocytes (set I) cultured in 2% FBS.

### Experimental set-up for astrocyte multi-omics Reactive vs control

Day 1 (fully differentiated) BIONi037-A or Kolf2.1J astrocytes were plated at 17K cells/cm2 in AM + 2% FBS in uncoated 10 cm dishes. At day 2, medium was replaced for AM without FBS after one PBS wash. At day 3, medium was replaced for AM without FBS supplemented with a reactive cytokine cocktail 30 ng/ml TNF (300-01A, Peprotech), 3 ng/ml Il-1α (AF-200-01A, Peprotech) and 400 ng/ml C1q (204876, Sigma Aldrich) or AM without FBS supplemented with an equal amount of PBS + 0,1% BSA (Tebu-Bio, 1501) as control. After 24h, iAstrocytes were collected by accutase dissociation after one PBS wash. iAstrocytes from each replicate dish were divided over 2 vials in which 1 million (Kolf2.1J) or 500K iAstrocytes (BIONi037-A) were pelleted for proteomic and lipidomic analysis. Cell pellets were stored at −80°C before further processing.

For iAstrocytes on the WTC11 background, differentiation was performed as previously described^90^, with minor modifications. Briefly, WTC11 iPSCs were edited to introduce a doxycycline-induced cassette driving pro-astrocyte transcription factors NFIA and SOX9. These iPSCs were differentiated into neural precursor cells (NPCs) using dual-SMAD inhibition/embroid body formation. NPCs were purified by fluorescence-activated cell sorting for CD133+/CD271– populations. Purified NPCs were further differentiated into iAstrocytes by doxycycline treatment (2 μg/ml, Millipore Sigma, D9891) and exposure to Astrocyte Medium (AM) (ScienCell, 1801) for 20 days. For experiments using serum-containing growth conditions, Day 20 iAstrocytes were plated at 20k cells/cm2 in phenol red-free AM (prfAM) (ScienCell, 1801-prf) overnight, with a full media change to fresh prfAM on Day 21. Media was changed to fresh prfAM every 2 days. On Day 25, cells were treated with 30ng/ml TNF (300-01A, Peprotech), 3nl/ml il-1α (AF-200-01A, Peprotech) and 400 ng/ml C1q (204876, Sigma Aldrich). After 24h media was removed, cells were washed with 1X DPBS and cell pellets collected and stored at −80°C before further processing for lipidomics. Studies with human iPSCs at UCSF were approved by the Human Gamete, Embryo and Stem Cell Research Committee. Informed consent was obtained from the human subjects when the iPSC lines were originally derived.

### Lipidomics of iPSC-derived TMEM106B KO neurons

TMEM106B KO iPSCs, genetically engineered from the parental KOLF2.1J iPSC line^54^, were obtained from the iPSC Neurodegenerative Disease Initiative (iNDI)^55^ via the Jackson Laboratory (JAX). These iPSCs, along with wild-type parental KOLF2.1J iPSCs, were cultured in feeder free conditions on Matrigel in E8 media (Life Technologies), and passaged via accuatase dissociation followed by E8 plus Chroman-I rock inhibitor. A piggybac-based tet-on NGN2 transgene cassette was stably integrated into the genome of iPSCs as described (https://www.protocols.io/view/indi-piggybac-to-hngn2-transfection-protocol-versi-q26g744b1gwz/v1) followed by puromycin selection to eliminate iPSCs that did not successfully integrate the transgene. iNeurons were differentiated on 6 well poly-ornithine coated dishes, with a plating density of 500,000 cells/well at d4, via doxycycline-induced expression of the NGN2 transgene, as described in (https://www.protocols.io/view/indi-transcription-factor-ngn2-differentiation-of-b2whqfb6.html). Cells were harvested at d21 post dox addition and snap frozen prior to lipidoic analysis. Cells from two wells were combined into one 1.5-mL tube as one sample pellet. Experiments were done as three independent replicates with 3 samples per replicate.

### Sample preparation for proteomics analysis

Frozen pellets corresponding to ∼500k cells were dissolved in 25 μl of PBS supplemented with 1 tab of cOmplete™, Mini, EDTA-free Protease Inhibitor per 50 ml. One volume equivalent of 2× lysis buffer (100 mM HEPES pH 8.0, 50 mM DTT, 4% (w/v) SDS) was added. Samples were sonicated in a Bioruptor Plus (Diagenode, Belgium) for 10 cycles with 1 min ON and 30 s OFF with high intensity at 20 °C. Samples were heated for 5 min at 95°C and a second sonication cycle was performed as described above. Samples were alkylated using freshly made 15 mM iodoacetamide (IAA) (Sigma-Aldrich #I1149) for 30 min at room temperature in the dark. Subsequently, proteins were acetone precipitated and digested using LysC (PTMScan, Cell signaling, #39003) and trypsin (Promega sequencing grade #V5111), as described by^91^. The digested proteins were then acidified with 10% (v/v) trifluoracetic acid and desalted using Waters Oasis® HLB µElution Plate 30 µm (Waters #186001828BA) following manufacturer instructions. The eluates were dried down using a vacuum concentrator and reconstituted in 5% (v/v) acetonitrile, 0.1% (v/v) formic acid. Samples were transferred to an MS vial, diluted to a concentration of 1 µg/µl, and spiked with iRT kit peptides (Biognosys AG #Ki-3002-2) prior to analysis by LC-MS/MS.

### Proteomics data acquisition

Peptides were separated in trap/elute mode using the nanoAcquity MClass Ultra-High Performance Liquid Chromatography system (Waters, MA, USA) equipped with trapping (nanoAcquity Symmetry C18, 5 μm, 180 μm × 20 mm) and an analytical column (nanoAcquity BEH C18, 1.7 μm, 75 μm × 250 mm). Solvent A was water and 0.1% formic acid, and solvent B was acetonitrile and 0.1% formic acid. 1 μl of the samples (∼1 μg on column) were loaded with a constant flow of solvent A at 5 μl/min onto the trapping column. Trapping time was 6 min. Peptides were eluted via the analytical column with a constant flow of 0.3 μl/min. During the elution, the percentage of solvent B increased nonlinearly from 0–40% in 120 min. The total run time was 145 min, including equilibration and conditioning. The LC was coupled to an Orbitrap Exploris 480 (Thermo Fisher Scientific, Germany) using the Proxeon nanospray source. The peptides were introduced into the mass spectrometer via a Pico-Tip Emitter 360 μm outer diameter × 20 μm inner diameter, 10-μm tip (New Objective) heated at 300 °C, and a spray voltage of 2.2 kV was applied. The capillary temperature was set at 300°C. The radio frequency ion funnel was set to 30%. For DIA data acquisition, full scan mass spectrometry (MS) spectra with a mass range 350–1650 m/z were acquired in profile mode in the Orbitrap with the resolution of 120,000 FWHM. The default charge state was set to 3+. The filling time was set at a maximum of 60 ms with a limitation of 3 × 10^6^ ions. DIA scans were acquired with 40 mass window segments of differing widths across the MS1 mass range. Higher collisional dissociation fragmentation (stepped normalized collision energy; 25, 27.5, and 30%) was applied, and MS/MS spectra were acquired with a resolution of 30,000 FWHM with a fixed first mass of 200 m/z after accumulation of 3 × 10^6^ ions or after filling time of 35 ms (whichever occurred first). Data were acquired in profile mode. For data acquisition and processing of the raw data, Xcalibur 4.3 (Thermo Fisher Scientific, Germany) and Tune version 2.0 were used.

### Proteomics data analysis

DIA raw data were analyzed using the directDIA pipeline in Spectronaut v.18 (Biognosys, Switzerland) with BGS settings besides the following parameters: Protein LFQ method= QUANT 2.0, Proteotypicity Filter = Only protein group specific, Major Group Quantity = Median peptide quantity, Minor Group Quantity = Median precursor quantity, Data Filtering = Qvalue, Normalizing strategy = Local Normalization. The data were searched against a UniProt (Homo Sapiens, 20,375 entries) and a contaminants (247 entries) database. The identifications were filtered to satisfy FDR of 1 % on peptide and protein level. Relative protein quantification was performed in Spectronaut using a pairwise t-test performed at the precursor level followed by multiple testing correction according to^92^.

### RNA-seq analysis

RNA isolation, QC, preprocessing, and data analysis were performed as previously described from frozen pellets^93^. Briefly, total RNA was isolated from each sample using the Qiagen RNeasy mini kit. RNA samples for each subject were entered into an electronic tracking system and processed at the UCI Genomics Research and Technology Hub (GRTH). RNA was QCed using an Agilent Bioanalyzer and quantified by Nanodrop. RNA quality is measured as RIN values (RNA Integrity Number), and 260/280 and 260/230 ratios to evaluate any potential contamination. Only samples with RIN >8 are used for library prep and sequencing. Library prep processing was initiated with total RNA of 1 µg using a Ribo-Zero Gold rRNA depletion and Truseq Stranded total RNA kit. RNA was chemically fragmented and subjected to reverse transcription, end repair, phosphorylation, A-tailing, ligation of barcoded sequencing adapters, and enrichment of adapter-ligated cDNAs. RNA-seq libraries were titrated by qPCR (Kapa), normalized according to size (Agilent Bioanalyzer 2100 High Sensitivity chip). Each cDNA library was then subjected to Illumina (Novaseq 6000) paired end (PE), 100 cycle sequencing to obtain approximately 50-65M PE reads. Fastq were subject to QC and reads with quality scores (>Q15) collected. Raw reads were mapped to the GRCh38 reference genome using Hisat2 (v.2.2.1), QCed, and normalization and transformation before further exploratory and differential expression analysis. Raw counts were normalized and transformed using the ‘*regularized log’ transformation* pipeline from the *R* package *DESeq2*. Statistical analyses was performed in *R* and differentially expressed genes detected for each covariate using false discovery rate or Bonferroni adjustment for multiple testing correction. Principal component analysis (PCA) was performed using *plotPCA* function in R with default settings. Following regularized log transformation in DESeq2, the top 500 highly variable genes (HVGs) were used as input for PCA and clustering of samples. DESeq2 was used to assess the statistical difference between the ApoE genotypes. Subsequently, we used the differentially expressed genes for each comparison to perform gene set enrichment analyses using Webgestalt^94^.

### RNA-seq data comparison

Expression data of iPSC derived astrocytes from TCW *et al*. (2022) and Lin *et al*. (2018) were analyzed and downloaded using GEO2R (Barret et al., 2013). TCW *et al*. performed bulk RNA sequencing on four isogenic sets of APOE3/3 and APOE4/4 iPSC-derived astrocytes, as well as bulk RNA-seq on seven APOE3/3 and six APOE4/4 population iPSC-derived astrocyte lines. Lin *et al*. performed bulk RNA seq on one isogenic set of APOE3/3 and APOE4/4 iPSC-derived astrocytes. From all five isogenic sets and the population model we gained the differential gene expression data of APOE4/4 versus APOE3/3. We calculated the average log2FC in ApoE4 vs ApoE3 of the pathways that were the top 10 up- and downregulated pathways in our APOE4/4 vs APOE3/3 transcriptomics for each line or the population data. We compared the directionality in all the lines and further explored the gene expression of the pathways that had the same directionality for all the comparisons.

### Experimental set up baseline experiments

At day 1, iAstrocytes were plated at 30-40K cells/cm^2^ in 6-,12-,96-well plates depending on the specific experiment in AM + 2% FBS (ScienCell). 24h after plating (day 2) medium was exchanged for AM without FBS after one PBS wash. On day 3, if needed, medium was collected and stored at −20°C until further analysis by MSD cytokine ELISA. Attached iAstrocytes were either fixed by 3.7% formaldehyde (FA; Electron Microscopy Sciences, 15681) for 10-15 minutes at RT and stored at 4°C in 1X PBS for immunofluorescent staining, collected by accutase dissociation for flow cytometry, lysed in Laemmli sample buffer with DTT (LSB; made in-house) for western blot or lysed in RIPA buffer (made in-house) for BCA analysis.

### Experimental set up drug treatment experiments

At day 1, iAstrocytes were plated at 30-40K cells/cm^2^ in 6-,12-,96-well plates depending on the specific experiment in AM + 2% FBS (ScienCell). 24h after plating (day 2) medium was exchanged for AM without FBS after one PBS wash. At day 3, iAstrocytes were treated with 50µM cholesterol (C4951, Sigma Aldrich), 0.5 µM Avasimibe (PZ0190, Sigma Aldrich), the reactive cytokine cocktail, 30ng/ml TNF (300-01A, Peprotech), 3ng/ml Il-1α (AF-200-01A, Peprotech) and 400ng/ml C1q (204876, Sigma Aldrich) (dose 1), or lower titrated doses of the cocktail indicated by 0.5 (resp. 15ng/ml TNF, 1.5ng/ml Il-1α and 200ng/ml C1Q), 0.25, 0.125 etc. Where indicated iAstrocytes were pre-incubated for 1h with 0.5 µM Avasimibe (PZ0190, Sigma Aldrich), 10µM Cholesterol (C4951, Sigma Aldrich) or 0.5µM Atorvastatin (HY-17379, MedChemExpress) before combined incubation with one of the previously mentioned treatments. 24h later at day 4, if needed, medium was collected and stored at −20°C until further analysis by MSD cytokine ELISA. Attached iAstrocytes were either fixed by 3.7% FA for 10-15 minutes at RT and stored at 4°C in 1X PBS for immunofluorescent staining, collected by accutase dissociation for flow cytometry, lysed in LSB for western blot or lysed in RIPA buffer for BCA analysis.

### Mesoscale discovery (MSD) cytokine measurements

Medium was thawed and cellular debris was removed by a 5 min spin at 2000g. Cytokine levels were determined by MSD Il-6 V-plex (K151QXD-2, MSD) according to manufacturer’s protocol. Medium samples were analyzed either undiluted or diluted 1/5 or 1/10 times in diluent 2 when iAstrocytes were treated with the reactive cocktail. Raw cytokine values per well were normalized over nuclei number per well based on fluorescent staining of the fixed iAstrocytes in the plate or over protein content per well determined by Pierce™ BCA protein assay kit (Thermo Scientific, 23225). Pierce™ BCA protein assay was performed in a microplate as described in the user guide provided by Thermo Scientific.

### Immunofluorescent stainings and imaging

After fixation, iAstrocytes were permeabilized with 0.5% Triton X-100 (Fisher Scientific, T/3751/08) for 5 min at RT and blocked in PBS with 0.1% Triton X-100 and 2% NGS (Fisher Scientific, 11540526) for 30 minutes at RT. Next, the iAstrocytes were incubated with the primary antibodies in blocking solution for 2 hours at RT or overnight at 4°C. The following primary antibodies were used: anti-perilipin 2 (15294-1-AP, Proteintech), anti-AQP4 (AQP-004, Alomone labs), anti-GFAP (173 004, Synaptic systems), anti-human HLA-A,B,C Antibody (311406, BioLegend). After 3 washes in 1X PBS, the iAstrocytes were incubated with Alexa Fluor secondary antibodies (Invitrogen; 1:1000), combined with DAPI (Carl Roth, 6843.1) and optional Lipidspot 488 (70065, Bio Connect; 1:1000) in blocking solution for 1 hour at RT. The iAstrocytes were washed 3 times in 1X PBS and either left in PBS to be imaged on the CellInsight CX7 LED Pro HCS Platform (Fisher Scientific, Hampton, NH, USA) or mounted on coverslips with Mowiol® 4-88 (Sigma Aldrich, 475904) for confocal imaging on a Nikon Ti-Eclipse microscope, equipped with a confocal scanner model A1R+, using a 40× oil immersion objective (NA = 1.3). Image analysis was done using Columbus® version 2.5.2 (PerkinElmer, Waltham, MA, USA) after imaging on CX7 and using Fiji (Schindelin et al., 2012) after confocal imaging on the Nikon Ti-Eclipse.

### Flow cytometry

After accutase dissocation, FACS buffer, DPBS + 2% FBS (Fisher Scientific, A5256701), was added to a volume of max. 300 µL and the iAstrocytes were transferred to a round bottom 96 well plate. After centrifuging for 1 minute at 2000 RPM, the iAstrocytes were stained with 1:50 PE anti-human HLA-A,B,C Antibody (BioLegend, 311406) and DAPI for 30 minutes at 4°C. Following another centrifugation step, the iAstrocytes were fixed with 2% formaldehyde (Sigma Aldrich, P6148) for 15 minutes at RT. After a last centrifugation step, the iAstrocytes were transferred to FACS tubes and 10.000 iAstrocytes of each sample were analyzed using BD LSRFortessa X-20 (BD Biosciences, Franklin Lakes, NJ, USA). Data analysis was done in FlowJo™ (BD Biosciences, Franklin Lakes, NJ, USA). First, a life gate was set using the forward and side scatter area, after which the single cells were gated using the forward scatter area and height. The geometric mean of the fluorescent intensity was used in the analyses.

### Western blot

After lysing the iAstrocytes with LSB, samples were denatured at 95°C for 5 min. The samples were shortly vortexed and loaded onto 4-15% Criterion TGX Stain-Free gel (BIO-RAD, 5678085). After running the gel (90V, 30 min followed by 150V, 45 min) the gel was transferred to a LF PVDF membrane using the Trans-Blot Turbo RTA Midi 0.45 µm LF PVDF Transfer Kit (BIO-RAD, 1704275). After blocking in 5% skim milk powder (Sigma Aldrich, 115363) for 1h at RT on a shaker, the membrane was incubated with antibodies against HLA Class I Heavy Chain (in-house NKI, 1:250) and GAPDH (elabscience, E-AB-40337, 1:3000) overnight at 4°C on a rocking plate. The following day the membrane was incubated with secondary antibodies IRDye® 800CW or IRDye® 680RD (LI-COR, 1:10000) for 1h at RT on a shaker. Afterwards, the membrane was scanned using the LI-COR® Odyssey® Fc Imaging System (LI-COR, Cambridge, UK). Analysis was done using Image Studio™ Lite 5.2.5 Software (LI-COR, Cambridge, UK) by calculating the median intensity of the bands minus the background above and below the bands.

### Statistical analysis

Statistical analysis was performed in Graphpad Prism version 10.2.3 (GraphPad Software, Boston, MA, USA) or Rstudio version 2023.9.1 (RStudio, PBC, Boston, MA, USA). The statistical tests used and the number of n (sample size) are annotated in the figures.

## Data availability

All RNAseq and proteomics data generated during this study will be deposited upon publication and are available upon request from the corresponding author.

## Acknowledgements

We thank members of the Van der Kant lab, Kim de Kleijn and Matthijs Verhage for helpful discussions of this work. We thank Ruud Wijdeven and the Advanced Cytometry Research Core Facility of the VU University Medical Center (O2Flow) for technical support. We thank Jie Wu for assistance in processing RNAseq data. We also thank the Genomics Research and Technology Hub Shared Resource of the Cancer Center Grant (CA-62203) at the University of California Irvine for facilities and carrying out RNA sequencing.

## Author contributions

FMF designed, performed and analyzed experiments on all aspects of the study

SJK designed, performed and analyzed experiments on all aspects of the study

LW performed lipidomics experiments and co-developed the Neurolipid Atlas tool

RD co-developed the Neurolipid Atlas tool

IK performed experiments to generate astrocyte multi-omics dataset

LE performed experiment to generate microglia lipidomics dataset

RM provided analysis for RNAseq

JH performed experiment to generate neuron lipidomics dataset

NP co-designed and performed proteomics experiments

NB performed lipidomic measurements

DOJ developed code for the Neurolipid Atlas

LEJ developed code for the Neurolipid Atlas

AJC provided pilot data, contributed

ALD data and co-developed the Neurolipid Atlas

CET provided human brain samples for lipidomics sampling

AG generated ALS neurons for lipidomics

IVLR designed and performed reactive astrocyte lipidomics experiments from WTC11 line

HY generated generated TMEM line neurons for lipidomics

MW generated lipidomics data on TMEM lines

AMI designed and oversaw generation of ALS lipidomics data

MK designed and oversaw generation reactive astrocyte data on WTC11 line

DKV designed and oversaw generation of microglia lipidomics data

TL provided human brain samples for lipidomics sampling

LMT co-designed multi-omics experiments and provided RNAseq analysis

AO co-designed multi-omics experiments and analyzed proteomics data

YM wrote code required for the Neurolipid Atlas and analysis of multi-omics data and oversaw the development of the original iSODA software tool

MG co-developed the project, oversaw all experiments and lipidomics analysis and co-developed Neurolipid Atlas

RK co-developed the project, oversaw all experiments, multi-omics analysis and iPSC experiments and co-developed Neurolipid Atlas

## Supplementary figures

**Supplementary Figure 1.**
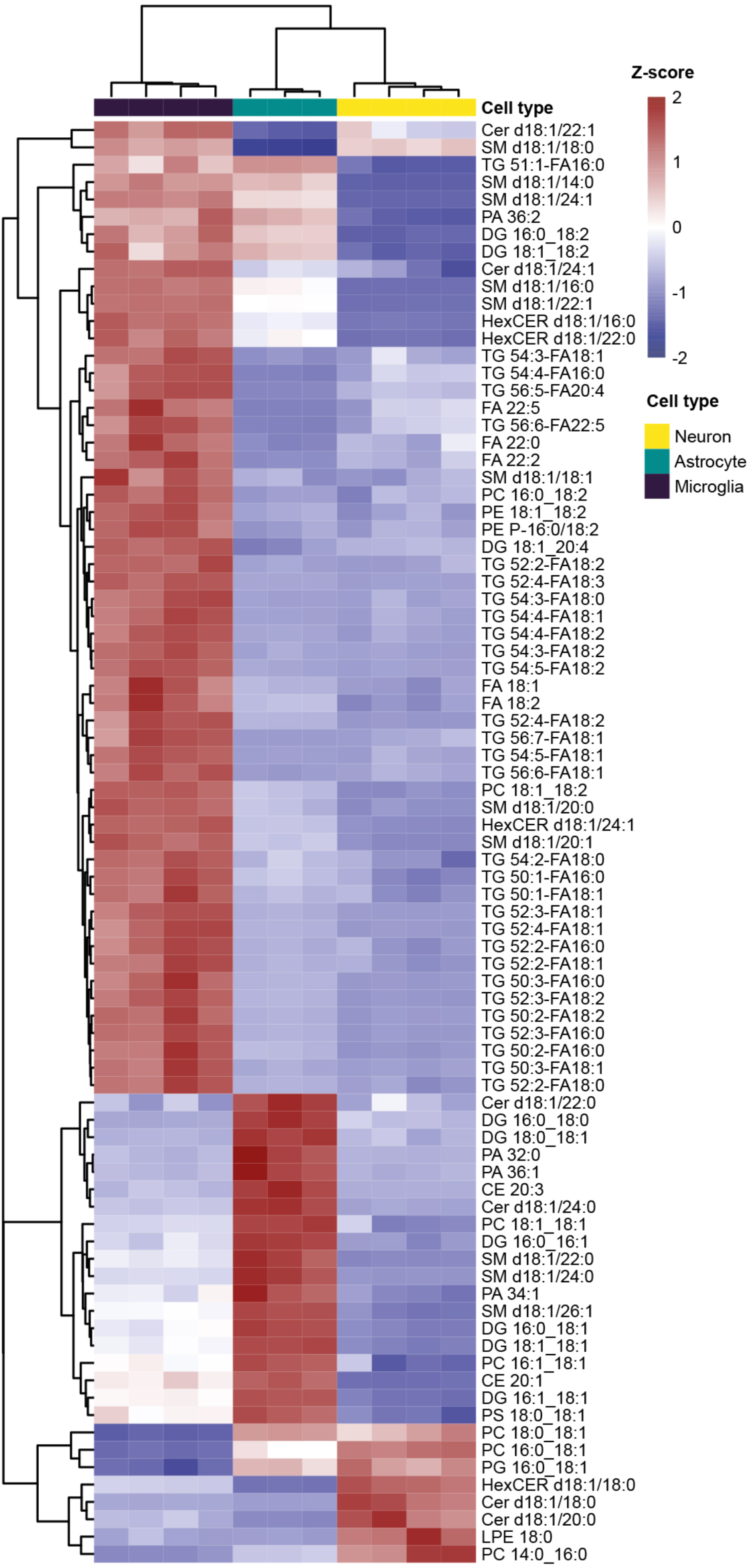
Lipotypes (most differentiating species) of human iPSC-derived neurons, astrocytes and microglia. A) Heatmap of most differentiating lipid species different between iPSC-derived neurons, astrocytes and microglia. alpha = 0.8.

**Supplementary Figure 2.**
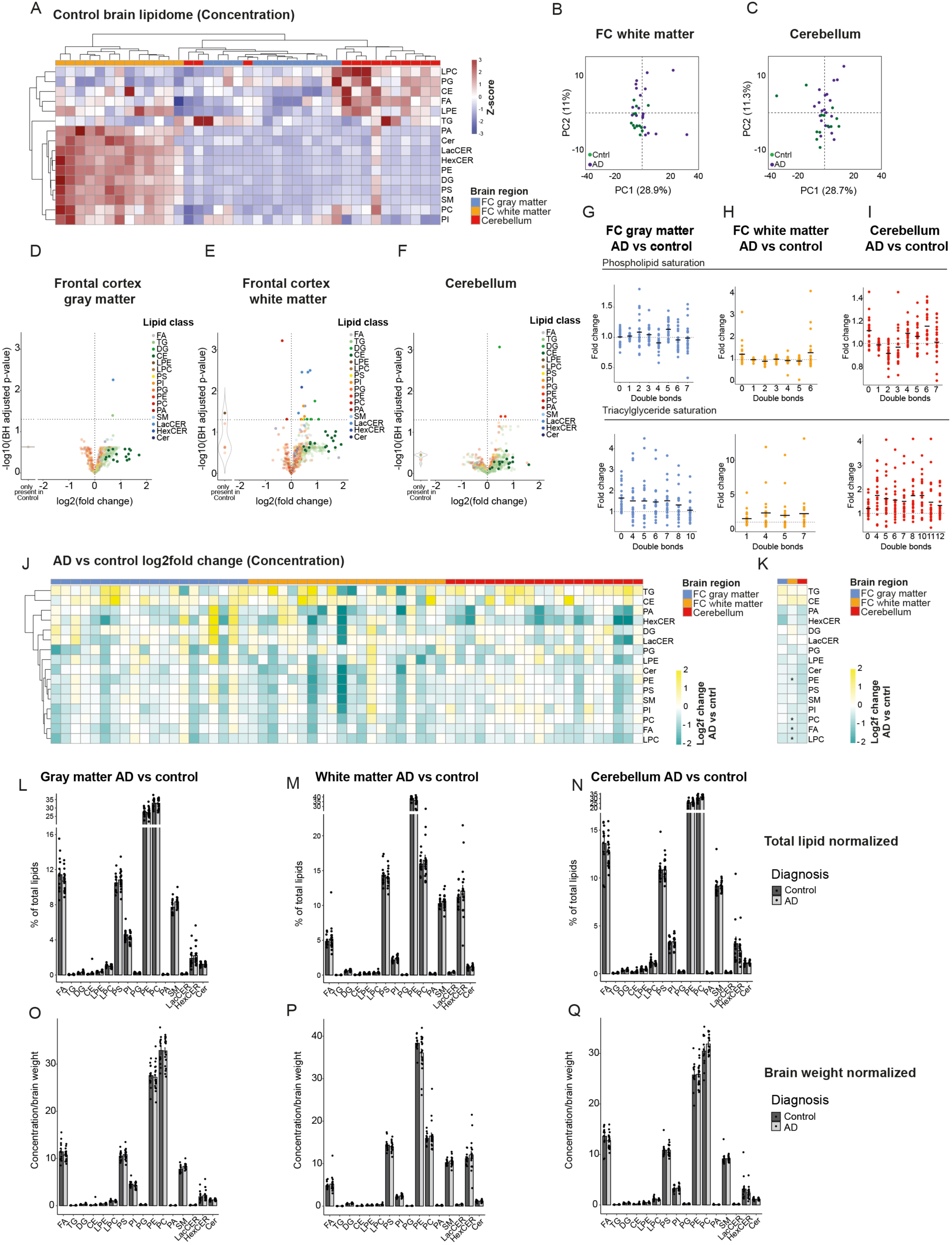
Extended analysis of human (AD) brain lipidomics. A) Heatmap with Z-scored lipid class concentrations (nmol/mg brain material) from indicated brain regions (control subjects only). B-C) PCA plot of unbiased lipidomic analysis of AD (purple) and control (green) brain tissue samples from (FC) white matter (B) and Cerebellum (C). D-F) Volcano plots of individual lipid species in AD vs control brain tissue for FC gray matter (D), FC white matter (E) and Cerebellum (F) as a % of total lipidome. CE species are highlighted. G-I) Fold change of phospholipid and TG species with indicated number of double bonds in AD vs control samples from FC gray matter (G), FC white matter (H) and Cerebellum (I) from % of total lipids. J) Heatmap depicting changes in lipid classes (concentrations) for individual AD samples compared to the average of control samples. Log2fold change plotted independent for each donor and each brain area. K) Average log2fold change in AD subject group compared to control samples per lipid class and brain area. Data from (J). L-N) Bar graphs present individual lipid class changes in FC gray matter (L), FC white matter (M) and Cerebellum (N) in AD versus control samples (group level) as % of total lipids. O-Q) Bar graphs present individual lipid class levels in FC gray matter (O), FC white matter (P) and Cerebellum (Q) in AD and control samples (group levels) as concentrations.

**Supplementary Figure 3.**
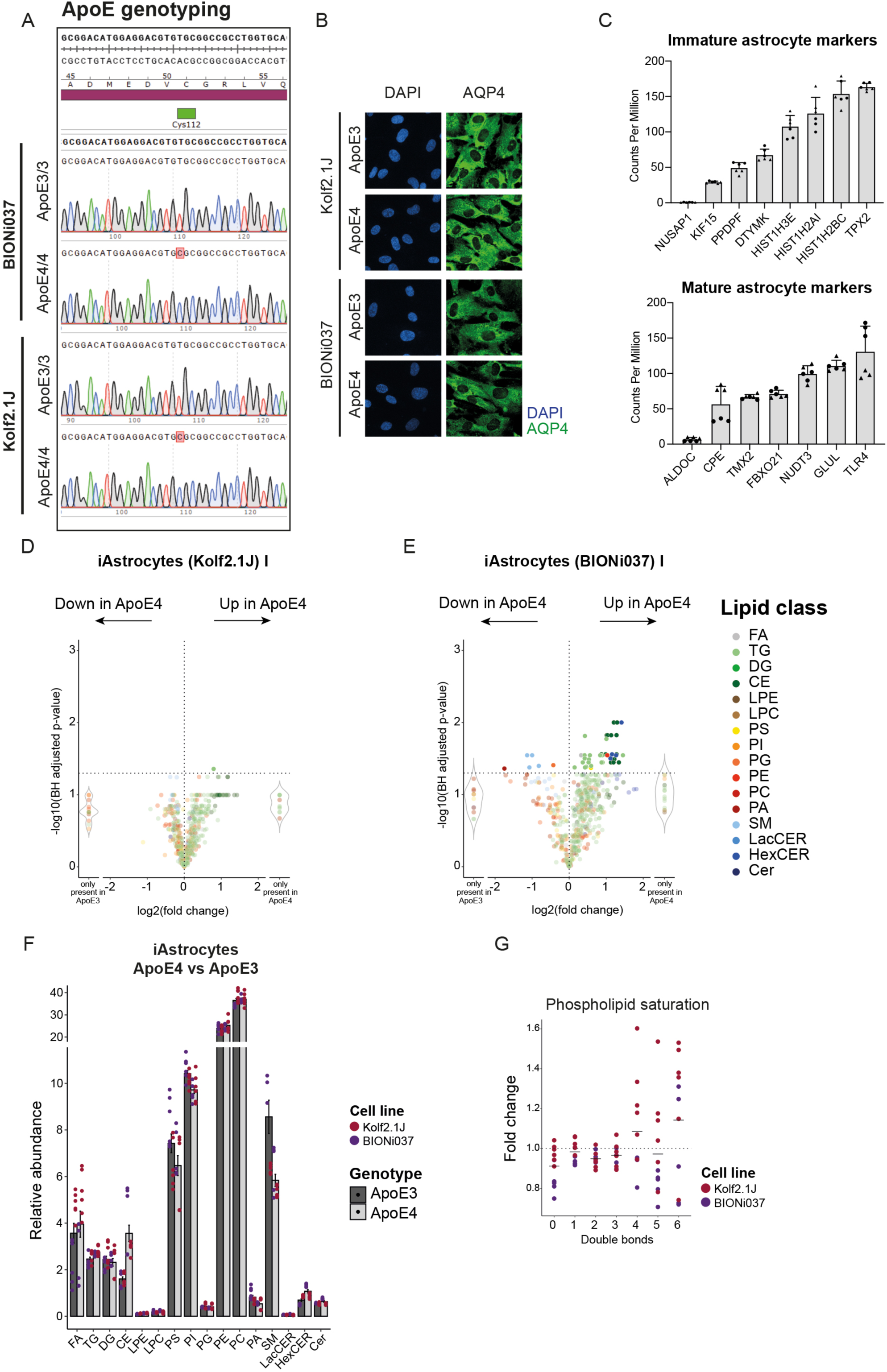
Extended lipidomic analysis of human isogenic APOE3/3 and APOE4/4 iPSC-derived astrocytes. A) Representative sequencing result for confirming cell identity and expected ApoE genotype B) Image of differentiated iAstrocytes, from both BIONi037 and Kolf2.1J lines. Scale bar = 25mm C) Gene expression levels (as determined by RNAseq) of indicated mature and immature astrocyte markers in our iPSC-derived astrocytes. n=6 wells BIONi037 (n=3 ApoE3 n=3 ApoE4) D-E) Volcano plots of lipid species in Kolf2.1J (D) or BIONi037 (E) ApoE4 vs ApoE3 iAstrocytes from a second independent lipidomics experiment. n=3 wells for Kolf2.1J, n=2 for BIONI037 (one BIONi037 ApoE4 sample was removed as outlier). F) Bar graphs of lipid classes in ApoE3 and ApoE4 iAstrocytes from BIONi037 and Kolf2.1J background (% of total). n=6 samples Kolf2.1J n=5 BIONI037 G) Fold change of phospholipid species with indicated number of double bonds in ApoE4 vs ApoE3 iAstrocytes.

**Supplementary Figure 4.**
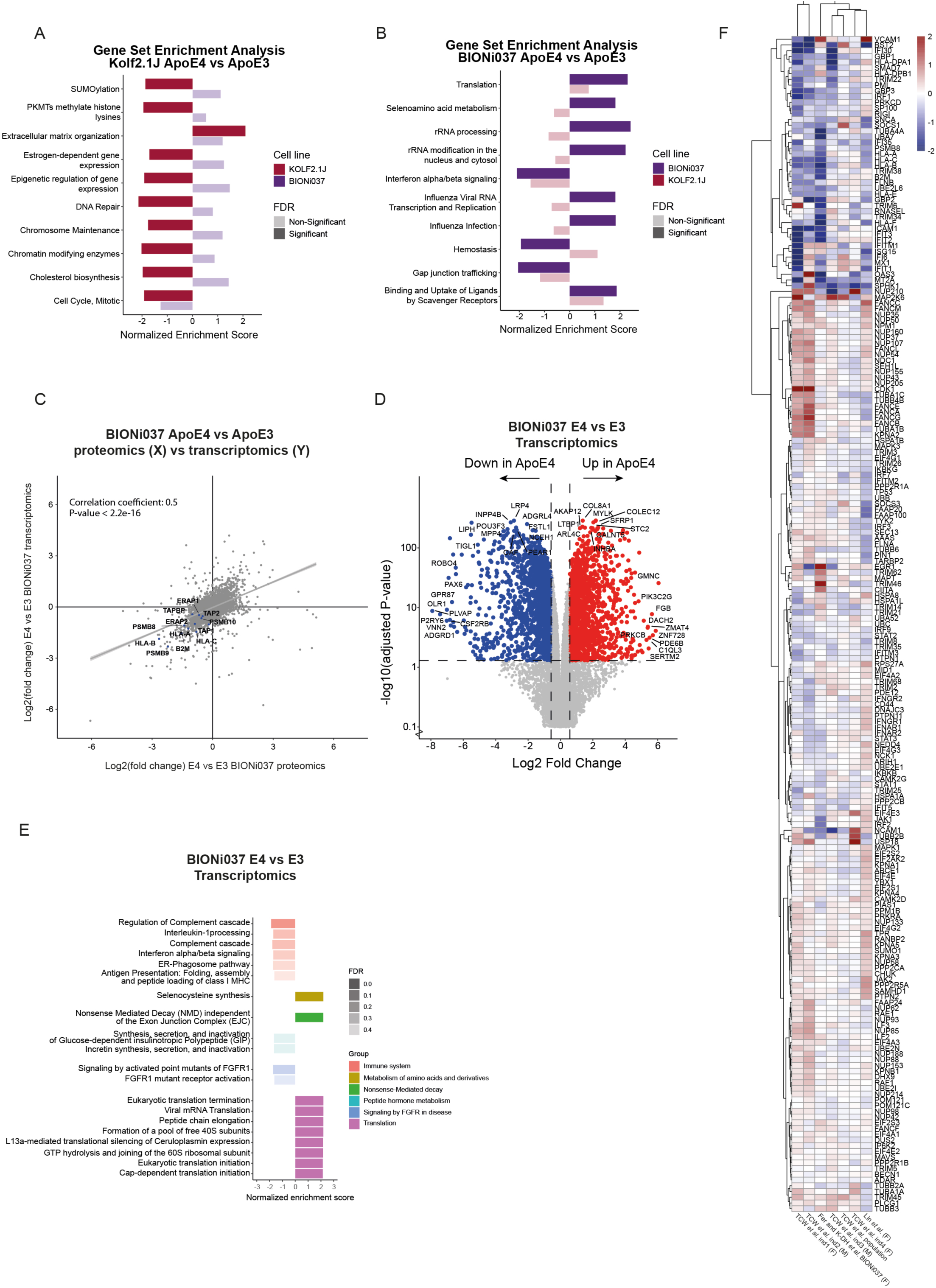
Extended proteomic and transcriptomic analysis of human isogenic APOE3/3 and APOE4/4 iPSC-derived astrocytes. A-B) Gene-set enrichment analysis was performed on proteomics data from ApoE4 versus ApoE3 iAstrocytes for the Kolf2.1J and the BIONi037 lines. Plotted are the enrichment scores (for both lines) for the reactome pathways significantly enriched in Kolf2.1J (A) and pathways significantly enriched in BIONI037 (B). C) Scatterplot of changes in protein levels (as measured by proteomics) versus changes in matching RNA expression (transcriptomics) in BIONi037 ApoE4 vs ApoE3 iAstrocytes. MHC-I and immunoproteasome pathway genes are indicated. D-F) Heatmaps showing changes of indicated genes in ApoE4 vs ApoE3 iAstrocytes from our study and previous studies (as indicated) within consistently altered pathways as identified by transcriptomics in Figure 4K. G) Transcriptomics data from ApoE4 vs ApoE3 BIONi037 iAstrocytes. Log2fold change of differentially expressed genes (DEGs) in ApoE4 vs ApoE3 astrocytes. Top ten genes with highest log2fold change and top ten genes with most significant P-value are labeled. n=3 wells per genotype. H) Top 10 reactome pathways upregulated or downregulated (with lowest FDR) in ApoE4 vs ApoE3 by gene-set enrichment analysis of transcriptomics data.

**Supplementary Figure 5.**
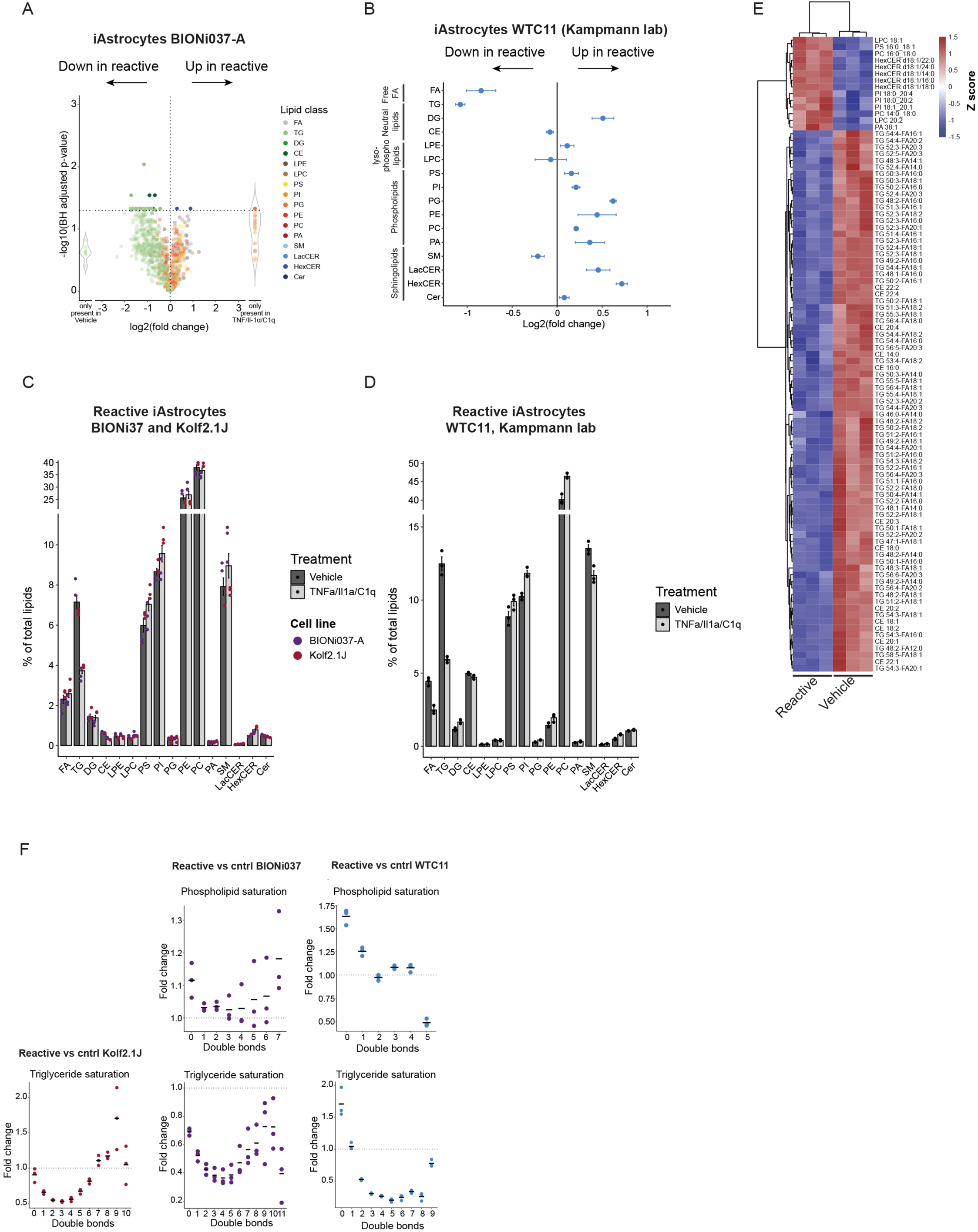
Extended lipidomics analysis of reactive human iPSC-derived astrocytes. A) Log2fold change of altered individual lipid species in reactive vs control iAstrocytes (BIONi037 ApoE3). n=3 wells reactive n=2 wells control. B) Summary data of changes in all detected lipid classes in reactive vs control iAstrocytes from the Kampmann lab (WTC11 iPSC line, ApoE3/3 cultured in 2% FBS). n=3 wells per condition. C-D) Bar graphs present changes in lipid classes in reactive vs control from Kolf2.1J and BIONi037 (C) or WTC11 (Kampmann lab) (D) iAstrocytes. Shown is % of total lipid, n=3 samples per line. E) Most differentiating lipid species in reactive vs control iAstrocytes (Kolf2.1J ApoE3) a=0.8. F) PL and TG saturation (number of double bonds) in Kolf2.1J, BIONi037 and WTC11 iAstrocytes.

**Supplementary Figure 6.**
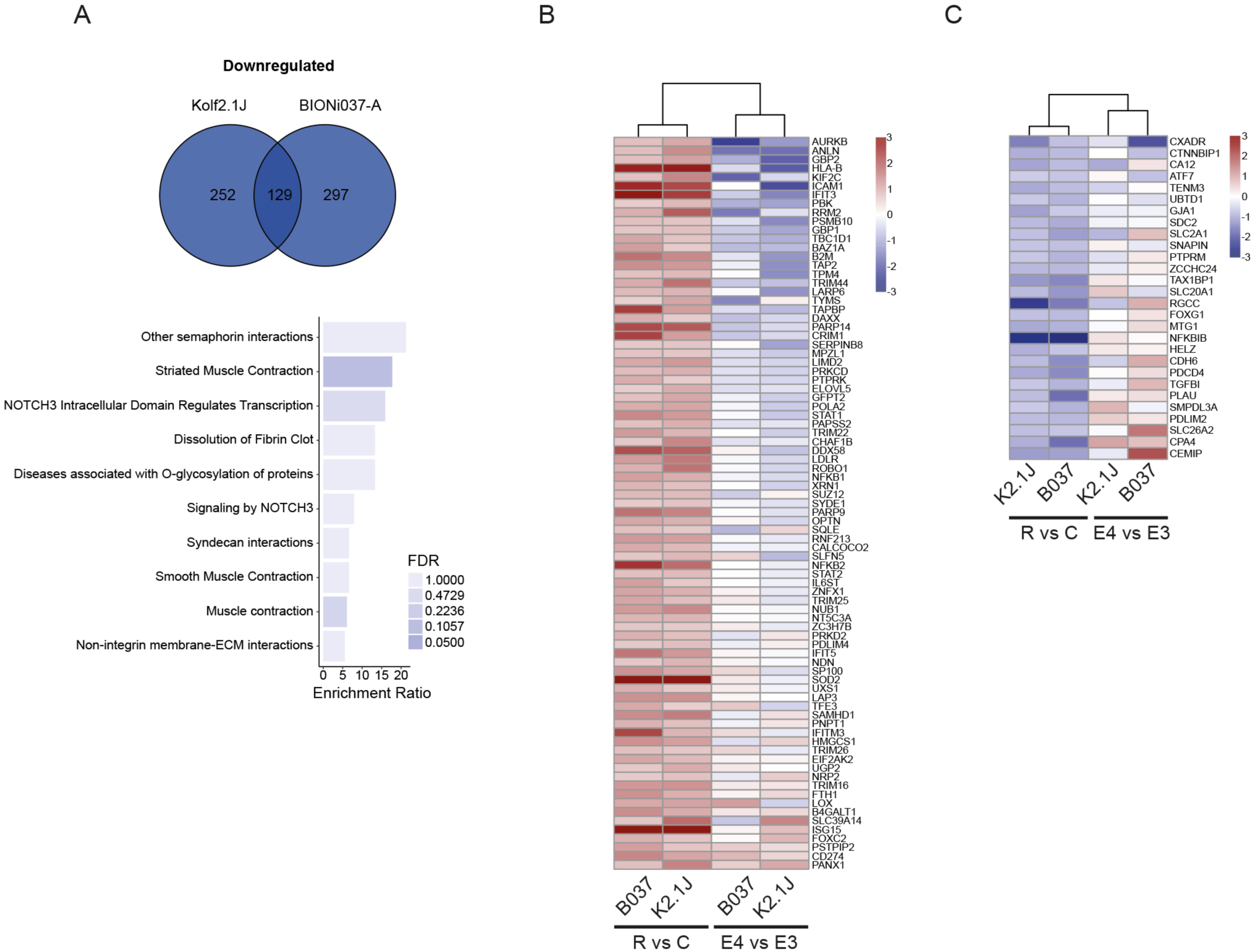
Extended proteomic analysis of reactive human iPSC-derived astrocytes. A) Venn diagram depicting the number of proteins that were significantly downregulated >1.25 fold (>0.3 log2fold) in reactive Kolf2.1J, BIONi037 and both iAstrocytes. Top 10 (non-significant) enriched reactome pathways detected by overrepresentation analysis are plotted. B) Heatmap shows the log2fold change of all proteins significantly upregulated >1.5 fold in both Kolf2.1J and BIONi037 reactive astrocytes. Next are the log2fold change values of these proteins in ApoE4 vs ApoE3 iAstrocytes. C) Heatmap shows the log2fold change of all proteins significantly downregulated >1.5 fold in Kolf2.1J and BIONi037 reactive astrocytes and the log2fold change values of these proteins in ApoE4 vs ApoE3 iAstrocytes.

**Supplementary Figure 7.**
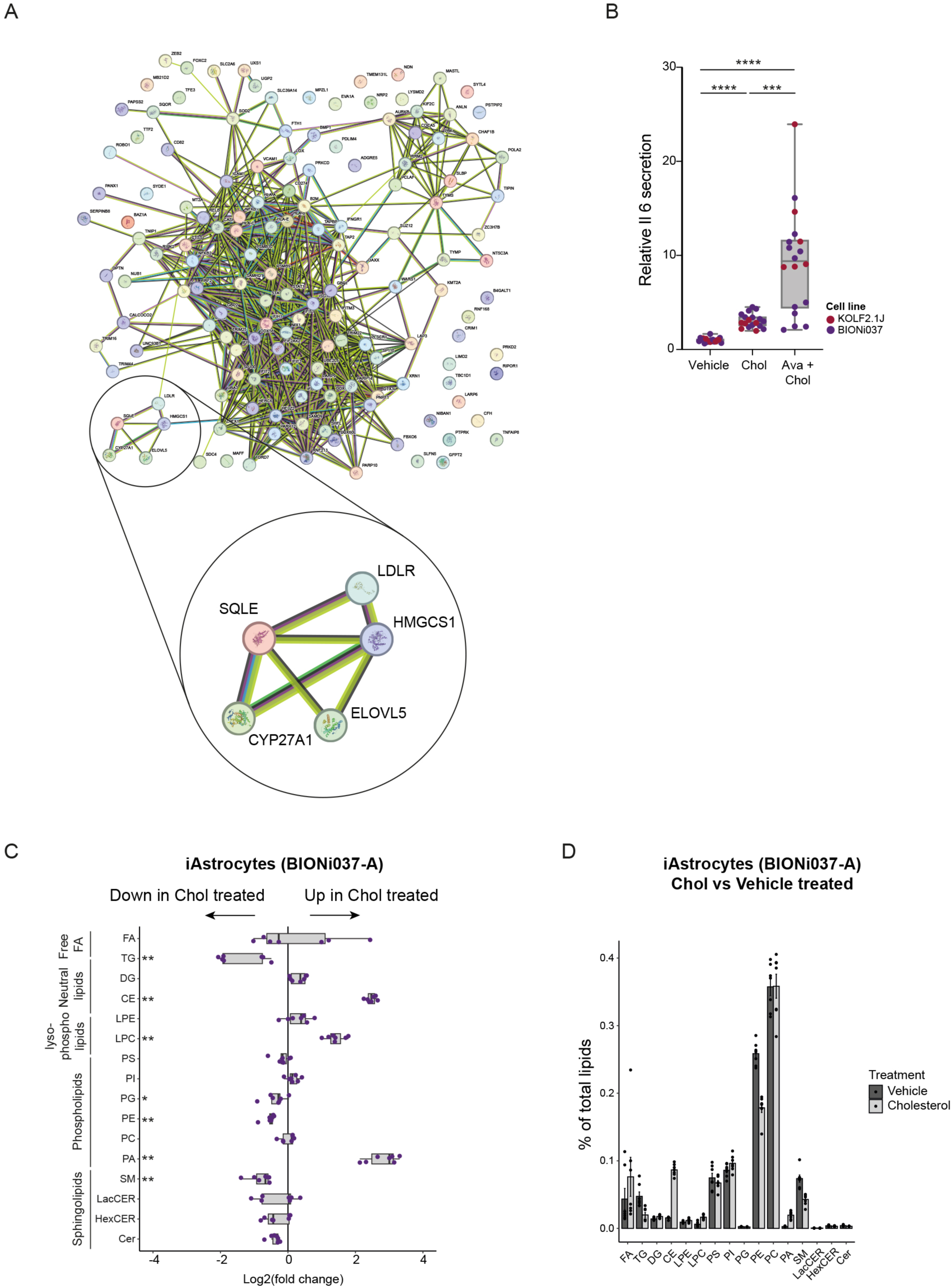
Cholesterol regulates astrocyte activation extended data. A) StringDB analysis of all proteins significantly upregulated >1.5 fold in Kolf2.1J and BIONi037 reactive astrocytes (dataset from figure 5). Zoom in shows cluster of genes related to cholesterol metabolism. B) Normalized Il-6 secretion in iAstrocytes (ApoE3), pre-treated with vehicle or avasimibe (0.5mM) for one hour and then treated for 24 hours with vehicle, cholesterol or cholesterol + avasimibe. n=6 (K2.1J) and n=12 (B037) from 4 independent experiments. ***P<0.001 Welch ANOVA with Dunnett’s multiple testing correction C) Summary data of changes in all detected lipid classes in cholesterol vs vehicle treated iAstrocytes (BIONi037-A). n=6 wells from 3 independent experiments. Mann-Whitney U test with Benjamini-Hochberg correction *=P<0.05. D) Bar graphs of lipid classes in cholesterol vs vehicle treated iAstrocytes from BIONi037-A background (% of total). n=6 wells from 3 independent experiments.

**Supplementary Table 1.**
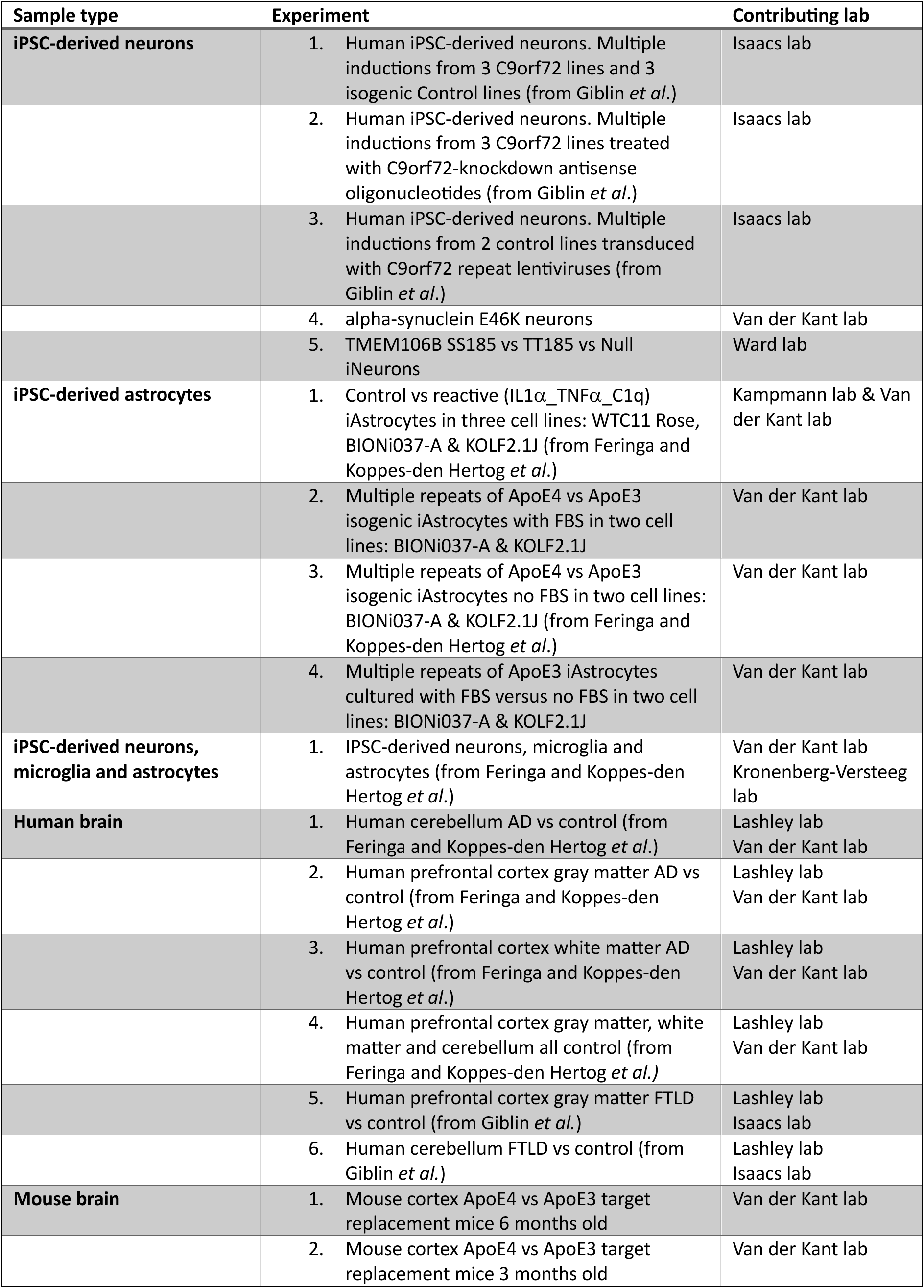
The Neurolipid Atlas: an open access data commons for brain lipid data. Description of datasets currently available on the Neurolipid Atlas.

## References

1. Yoon, J. H. et al. Brain lipidomics: From functional landscape to clinical significance. Sci. Adv. 8, 9317 (2022).

2. Kollmann, K. et al. Cell biology and function of neuronal ceroid lipofuscinosis-related proteins. Biochim. Biophys. Acta 1832, 1866–1881 (2013).

3. Vance, J. E. Lipid imbalance in the neurological disorder, Niemann-Pick C disease. FEBS Lett. 580, 5518–5524 (2006).

4. Chan, R. B. et al. Comparative Lipidomic Analysis of Mouse and Human Brain with Alzheimer Disease. J. Biol. Chem. 287, 2678–2688 (2012).

5. de Leeuw, S. M. et al. APOE2, E3, and E4 differentially modulate cellular homeostasis, cholesterol metabolism, and inflammatory response in isogenic iPSC-derived astrocytes. Stem Cell Reports 17, 110 (2022).

6. Di Paolo, G. & Kim, T. W. Linking lipids to Alzheimer’s disease: Cholesterol and beyond. Nature Reviews Neuroscience 12, 284–296 (2011).

7. Kant, R. van der et al. Cholesterol Metabolism Is a Druggable Axis that Independently Regulates Tau and Amyloid-β in iPSC-Derived Alzheimer’s Disease Neurons. Cell Stem Cell 24, 363 (2019).

8. Lin, Y. T. et al. APOE4 Causes Widespread Molecular and Cellular Alterations Associated with Alzheimer’s Disease Phenotypes in Human iPSC-Derived Brain Cell Types. Neuron 98, 1141–1154.e7 (2018).

9. Sienski, G. et al. APOE4 disrupts intracellular lipid homeostasis in human iPSC-derived glia. Sci. Transl. Med. 13, (2021).

10. TCW, J., et al. Cholesterol and matrisome pathways dysregulated in astrocytes and microglia. Cell 185, 2213–2233.e25 (2022).

11. Fanning, S. et al. Lipidomic Analysis of α-Synuclein Neurotoxicity Identifies Stearoyl CoA Desaturase as a Target for Parkinson Treatment. Mol. Cell 73, 1001–1014.e8 (2019).

12. Gedalya, T. Ben et al. α–Synuclein and PolyUnsaturated Fatty Acids Promote Clathrin Mediated Endocytosis and Synaptic Vesicle Recycling. Traffic 10, 218 (2009).

13. Boussicault, L. et al. CYP46A1 protects against NMDA-mediated excitotoxicity in Huntington’s disease: Analysis of lipid raft content. Biochimie 153, 70–79 (2018).

14. Boussicault, L. et al. CYP46A1, the rate-limiting enzyme for cholesterol degradation, is neuroprotective in Huntington’s disease. Brain 139, 953–970 (2016).

15. Leoni, V. & Caccia, C. The impairment of cholesterol metabolism in Huntington disease. Biochim. Biophys. Acta 1851, 1095–1105 (2015).

16. Nóbrega, C. et al. Restoring brain cholesterol turnover improves autophagy and has therapeutic potential in mouse models of spinocerebellar ataxia. Acta Neuropathol. 138, 837–858 (2019).

17. Chaves-Filho, A. B. et al. Alterations in lipid metabolism of spinal cord linked to amyotrophic lateral sclerosis. Sci. Rep. 9, (2019).

18. Giblin, A. et al. Neuronal polyunsaturated fatty acids are protective in FTD/ALS. bioRxiv 2024.01.16.575677 (2024).

19. Li, Y. et al. Microglial lipid droplet accumulation in tauopathy brain is regulated by neuronal AMPK. Cell Metab. (2024).

20. Liu, Y. & Wang, J. C9orf72-dependent lysosomal functions regulate epigenetic control of autophagy and lipid metabolism. Autophagy 15, 913–914 (2019).

21. Liu, Y. et al. A C9orf72-CARM1 axis regulates lipid metabolism under glucose starvation-induced nutrient stress. Genes Dev. 32, 1380–1397 (2018).

22. Valdez, C., Ysselstein, D., Young, T. J., Zheng, J. & Krainc, D. Progranulin mutations result in impaired processing of prosaposin and reduced glucocerebrosidase activity. Hum. Mol. Genet. 29, 716–726 (2020).

23. Marschallinger, J. et al. Lipid-droplet-accumulating microglia represent a dysfunctional and proinflammatory state in the aging brain. Nat. Neurosci. 23, 194–208 (2020).

24. Andreone, B. J. et al. Alzheimer’s-associated PLCγ2 is a signaling node required for both TREM2 function and the inflammatory response in human microglia. Nat. Neurosci. 23, 927– 938 (2020).

25. Haney, M. S. et al. APOE4/4 is linked to damaging lipid droplets in Alzheimer’s disease microglia. Nat. 2024 6288006 628, 154–161 (2024).

26. Nugent, A. A. et al. TREM2 Regulates Microglial Cholesterol Metabolism upon Chronic Phagocytic Challenge. Neuron 105, 837–854.e9 (2020).

27. Guttenplan, K. A. et al. Neurotoxic reactive astrocytes induce cell death via saturated lipids. Nat. 2021 5997883 599, 102–107 (2021).

28. Haynes, P. R. et al. A neuron–glia lipid metabolic cycle couples daily sleep to mitochondrial homeostasis. Nat. Neurosci. 27, 666 (2024).

29. Capolupo, L. et al. Sphingolipids control dermal fibroblast heterogeneity. Science (80-.). 376, (2022).

30. Bozek, K. et al. Organization and Evolution of Brain Lipidome Revealed by Large-Scale Analysis of Human, Chimpanzee, Macaque, and Mouse Tissues. Neuron 85, 695–702 (2015).

31. Byers, B. et al. SNCA Triplication Parkinson’s Patient’s iPSC-derived DA Neurons Accumulate α-Synuclein and Are Susceptible to Oxidative Stress. PLoS One 6, (2011).

32. Israel, M. A. et al. Probing sporadic and familial Alzheimer’s disease using induced pluripotent stem cells. Nature 482, 216–220 (2012).

33. Kondo, T. et al. Modeling Alzheimer’s disease with iPSCs reveals stress phenotypes associated with intracellular Aβ and differential drug responsiveness. Cell Stem Cell 12, 487–496 (2013).

34. Muratore, C. R. et al. The familial Alzheimer’s disease APPV717I mutation alters APP processing and Tau expression in iPSC-derived neurons. Hum. Mol. Genet. 23, 3523 (2014).

35. Haenseler, W. et al. A Highly Efficient Human Pluripotent Stem Cell Microglia Model Displays a Neuronal-Co-culture-Specific Expression Profile and Inflammatory Response. Stem Cell Reports 8, 1727–1742 (2017).

36. Handel, A. E. et al. Assessing similarity to primary tissue and cortical layer identity in induced pluripotent stem cell-derived cortical neurons through single-cell transcriptomics. Hum. Mol. Genet. 25, 989 (2016).

37. Nehme, R. et al. Combining NGN2 Programming with Developmental Patterning Generates Human Excitatory Neurons with NMDAR-Mediated Synaptic Transmission. Cell Rep. 23, 2509– 2523 (2018).

38. TCW, J., et al. An Efficient Platform for Astrocyte Differentiation from Human Induced Pluripotent Stem Cells. Stem cell reports 9, 600–614 (2017).

39. Nimsanor, N. et al. Generation of induced pluripotent stem cells derived from a 77-year-old healthy woman as control for age related diseases. Stem Cell Res. 17, 550–552 (2016).

40. Zhang, Y. et al. Rapid Single-Step Induction of Functional Neurons from Human Pluripotent Stem Cells. Neuron 78, 785–798 (2013).

41. Fong, L. K. et al. Full-length amyloid precursor protein regulates lipoprotein metabolism and amyloid-clearance in human astrocytes. J. Biol. Chem. 293, 11341–11357 (2018).

42. Ghorasaini, M. et al. Cross-Laboratory Standardization of Preclinical Lipidomics Using Differential Mobility Spectrometry and Multiple Reaction Monitoring. Anal. Chem. 93, 16369– 16378 (2021).

43. Su, B. et al. A DMS Shotgun Lipidomics Workflow Application to Facilitate High-throughput, Comprehensive Lipidomics. J. Am. Soc. Mass Spectrom. 32, 2655 (2021).

44. Fitzner, D. et al. Cell-Type- and Brain-Region-Resolved Mouse Brain Lipidome. Cell Rep. 32, (2020).

45. Ferris, H. A. et al. Loss of astrocyte cholesterol synthesis disrupts neuronal function and alters whole-body metabolism. Proc. Natl. Acad. Sci. U. S. A. 114, 1189–1194 (2017).

46. Pfrieger, F. W. & Ungerer, N. Cholesterol metabolism in neurons and astrocytes. Prog. Lipid Res. 50, 357–371 (2011).

47. Akyol, S. et al. Lipid profiling of Alzheimer’s disease brain highlights enrichment in glycerol(Phospho)lipid, and sphingolipid metabolism. Cells 10, 2591 (2021).

48. Bandaru, V. V. R. et al. ApoE4 disrupts sterol and sphingolipid metabolism in Alzheimer’s but not normal brain. Neurobiol. Aging 30, 591 (2009).

49. Raji, C. A., Lopez, O. L., Kuller, L. H., Carmichael, O. T. & Becker, J. T. Age, Alzheimer disease, and brain structure. Neurology 73, 1899 (2009).

50. Oshida, K. et al. Effects of Dietary Sphingomyelin on Central Nervous System Myelination in Developing Rats. Pediatr. Res. 2003 534 53, 589–593 (2003).

51. Corder, E. H. et al. Gene dose of apolipoprotein E type 4 allele and the risk of Alzheimer’s disease in late onset families. Science (80-.). 261, 921–923 (1993).

52. Farrer, L. A. et al. Effects of Age, Sex, and Ethnicity on the Association Between Apolipoprotein E Genotype and Alzheimer Disease: A Meta-analysis. JAMA 278, 1349–1356 (1997).

53. Fortea, J. et al. APOE4 homozygozity represents a distinct genetic form of Alzheimer’s disease. Nat. Med. 2024 1–8 (2024).

54. Pantazis, C. B. et al. A reference human induced pluripotent stem cell line for large-scale collaborative studies. Cell Stem Cell 29, 1685 (2022).

55. Ramos, D. M., Skarnes, W. C., Singleton, A. B., Cookson, M. R. & Ward, M. E. Tackling neurodegenerative diseases with genomic engineering: A new stem cell initiative from the NIH. Neuron 109, 1080 (2021).

56. Barbar, L. et al. CD49f Is a Novel Marker of Functional and Reactive Human iPSC-Derived Astrocytes. Neuron 107, 436–453.e12 (2020).

57. Giannisis, A. et al. Plasma apolipoprotein E levels in longitudinally followed patients with mild cognitive impairment and Alzheimer’s disease. Alzheimer’s Res. Ther. 2022 141 14, 1–17 (2022).

58. Martínez-Morillo, E. et al. Total apolipoprotein E levels and specific isoform composition in cerebrospinal fluid and plasma from Alzheimer’s disease patients and controls. Acta Neuropathol. 127, 633–643 (2014).

59. Arnaud, L. et al. APOE4 drives inflammation in human astrocytes via TAGLN3 repression and NF-κB activation. Cell Rep. 40, (2022).

60. Blumenfeld, J., Yip, O., Kim, M. J. & Huang, Y. Cell type-specific roles of APOE4 in Alzheimer disease. Nat. Rev. Neurosci. 25, 91 (2024).

61. Lee, S. et al. APOE4 drives transcriptional heterogeneity and maladaptive immunometabolic responses of astrocytes. doi:10.1101/2023.02.06.527204

62. Mhatre-Winters, I., Eid, A., Han, Y., Tieu, K. & Richardson, J. R. Sex and APOE Genotype Alter the Basal and Induced Inflammatory States of Primary Astrocytes from Humanized Targeted Replacement Mice. doi:10.1177/17590914221144549

63. Dakterzada, F. et al. Cerebrospinal fluid neutral lipids predict progression from mild cognitive impairment to Alzheimer’s disease. GeroScience 2023 1–14 (2023).

64. Hutter-Paier, B. et al. The ACAT Inhibitor CP-113,818 Markedly Reduces Amyloid Pathology in a Mouse Model of Alzheimer’s Disease. Neuron 44, 227–238 (2004).

65. Puglielli, L. et al. Acyl-coenzyme A: cholesterol acyltransferase modulates the generation of the amyloid β-peptide. Nat. Cell Biol. 3, 905–912 (2001).

66. Shibuya, Y. et al. Acyl-coenzyme A: CHolesterol acyltransferase 1 blockage enhances autophagy in the neurons of triple transgenic Alzheimer’s disease mouse and reduces human P301L-tau content at the presymptomatic stage. Neurobiol. Aging 36, 2248–2259 (2015).

67. Shibuya, Y., Chang, C. C. Y., Huang, L. H., Bryleva, E. Y. & Chang, T. Y. Inhibiting ACAT1/SOAT1 in microglia stimulates autophagy-mediated lysosomal proteolysis and increases Aβ1-42 clearance. J. Neurosci. 34, 14484–14501 (2014).

68. Blanchard, J. W. et al. APOE4 impairs myelination via cholesterol dysregulation in oligodendrocytes. Nat. 2022 6117937 611, 769–779 (2022).

69. Windham, I. A. et al. APOE traffics to astrocyte lipid droplets and modulates triglyceride saturation and droplet size. J. Cell Biol. 223, (2024).

70. Bryleva, E. Y. et al. ACAT1 gene ablation increases 24(S)-hydroxycholesterol content in the brain and ameliorates amyloid pathology in mice with AD. Proc. Natl. Acad. Sci. U. S. A. 107, 3081–3086 (2010).

71. Huttunen, H. J. et al. The Acyl-Coenzyme A: Cholesterol Acyltransferase Inhibitor CI-1011 Reverses Diffuse Brain Amyloid Pathology in Aged Amyloid Precursor Protein Transgenic Mice. J. Neuropathol. Exp. Neurol. 69, 777–788 (2010).

72. Huttunen, H. J. et al. Inhibition of acyl-coenzyme A: cholesterol acyl transferase modulates amyloid precursor protein trafficking in the early secretory pathway. FASEB J. 23, 3819–3828 (2009).

73. Murphy, S. R. et al. Acat1 knockdown gene therapy decreases amyloid-β in a mouse model of Alzheimer’s disease. Mol. Ther. 21, 1497–1506 (2013).

74. Meuwese, M. C. et al. ACAT inhibition and progression of carotid atherosclerosis in patients with familial hypercholesterolemia: the CAPTIVATE randomized trial. JAMA 301, 1131–1139 (2009).

75. Nissen, S. E. et al. Effect of ACAT Inhibition on the Progression of Coronary Atherosclerosis. N. Engl. J. Med. 354, 1253–1263 (2006).

76. Tardif, J. C. et al. Effects of the acyl coenzyme A:cholesterol acyltransferase inhibitor avasimibe on human atherosclerotic lesions. Circulation 110, 3372–3377 (2004).

77. Bhattarai, P. et al. Rare genetic variation in fibronectin 1 (FN1) protects against APOEε4 in Alzheimer’s disease. Acta Neuropathol. 147, 70 (2024).

78. Labib, D. et al. Proteomic Alterations and Novel Markers of Neurotoxic Reactive Astrocytes in Human Induced Pluripotent Stem Cell Models. Front. Mol. Neurosci. 15, 870085 (2022).

79. Akiyama, H. et al. Inflammation and Alzheimer’s disease. Neurobiol. Aging 21, 383–421 (2000).

80. Orre, M. et al. Reactive glia show increased immunoproteasome activity in Alzheimer’s disease. Brain 136, 1415–1431 (2013).

81. Overmyer, M. et al. Astrogliosis and the ApoE Genotype An Immunohistochemical Study of Postmortem Human Brain Tissue. Dement. Geriatr. Cogn. Disord. 10, 252–257 (1999).

82. Liu, C. C. et al. Cell-autonomous effects of APOE4 in restricting microglial response in brain homeostasis and Alzheimer’s disease. Nat. Immunol. 2023 2411 24, 1854–1866 (2023).

83. Yin, Z. et al. APOE4 impairs the microglial response in Alzheimer’s disease by inducing TGFβ-mediated checkpoints. Nat. Immunol. 2023 2411 24, 1839–1853 (2023).

84. Mancuso, R. et al. Xenografted human microglia display diverse transcriptomic states in response to Alzheimer’s disease-related amyloid-β pathology. Nat. Neurosci. 27, 886 (2024).

85. Claes, C. et al. The P522R protective variant of PLCG2 promotes the expression of antigen presentation genes by human microglia in an Alzheimer’s disease mouse model. Alzheimers. Dement. 18, 1765–1778 (2022).

86. Gazestani, V. et al. Early Alzheimer’s disease pathology in human cortex involves transient cell states. Cell 186, 4438–4453.e23 (2023).

87. Morana, O. et al. Identification of a New Cholesterol-Binding Site within the IFN-γ Receptor that is Required for Signal Transduction. Adv. Sci. 9, 2105170 (2022).

88. O’Carroll, S. M., Henkel, F. D. R. & O’Neill, L. A. J. Metabolic regulation of type I interferon production. Immunol. Rev. 323, (2024).

89. Shi, Y., Kirwan, P. & Livesey, F. J. Directed differentiation of human pluripotent stem cells to cerebral cortex neurons and neural networks. Nat. Protoc. 2012 710 7, 1836–1846 (2012).

90. Leng, K. et al. CRISPRi screens in human iPSC-derived astrocytes elucidate regulators of distinct inflammatory reactive states. Nat. Neurosci. 2022 2511 25, 1528–1542 (2022).

91. Buczak, K. et al. Spatially resolved analysis of FFPE tissue proteomes by quantitative mass spectrometry. Nat. Protoc. 2020 159 15, 2956–2979 (2020).

92. Storey, J. D. A direct approach to false discovery rates. J. R. Stat. Soc. Ser. B (Statistical Methodol. 64, 479–498 (2002).

93. Workman, M. J. et al. Large-scale differentiation of iPSC-derived motor neurons from ALS and control subjects. Neuron 111, 1191 (2023).

94. Liao, Y., Wang, J., Jaehnig, E. J., Shi, Z. & Zhang, B. WebGestalt 2019: gene set analysis toolkit with revamped UIs and APIs. Nucleic Acids Res. 47, W199–W205 (2019).

